# SRRF ‘n’ TIRF - FCS: Simultaneous spatiotemporal super-resolution microscopy

**DOI:** 10.1101/2020.02.26.965905

**Authors:** Jagadish Sankaran, Harikrushnan Balasubramanian, Wai Hoh Tang, Xue Wen Ng, Adrian Röllin, Thorsten Wohland

**Affiliations:** Department of Biological Sciences and NUS Centre for Bio-Imaging Sciences, National University of Singapore, 14 Science Drive 4, 117557 Singapore; Department of Statistics and Applied Probability, National University of Singapore, 6 Science Drive 2, 117546 Singapore; Department of Chemistry, National University of Singapore, 3 Science Drive 3, 117543 Singapore

**Keywords:** Super-resolution, Imaging fluorescence correlation spectroscopy, Number and brightness analysis, Super-resolution radial fluctuations, FCS diffusion law, Epidermal growth factor receptor

## Abstract

Super-resolution microscopy and single molecule fluorescence spectroscopy require mutually exclusive experimental strategies optimizing either time or spatial resolution. To achieve both, we implement a GPU-supported, camera-based measurement strategy that highly resolves spatial structures (~60 nm), temporal dynamics (≤ 2 ms), and molecular brightness from the exact same data set. We demonstrate the applicability and advantages of multi-parametric measurements to monitor the super-resolved structure and dynamics of two different biomolecules, the actin binding polypeptide LifeAct, and the epidermal growth factor receptor (EGFR). Simultaneous super-resolution of spatial and temporal details leads to an improved precision in estimating the diffusion coefficient of LifeAct in dependence of the cellular actin network. Multi-parametric analysis suggests that the domain partitioning of EGFR is primarily determined by EGFR-membrane interactions, possibly sub-resolution clustering and inter-EGFR interactions but is largely independent of EGFR-actin interactions. These results demonstrate that pixel-wise cross-correlation of parameters obtained from different techniques on the same data set enables robust physicochemical parameter estimation and provides new biological knowledge that cannot be obtained from sequential measurements.

## Introduction

Full knowledge of a biological system requires not only information on its spatial structure but also its temporal dynamics. However, the acquisition of structure and dynamics requires complementary, often mutually exclusive optimization strategies^1^. Spatial resolution depends on the number of photons collected and thus sets a lower limit on acquisition time. Molecular dynamics requires acquisition times shorter than the dynamics of interest and thus sets an upper limit. Since these two limits, in general, do not lead to an overlap region, the combination of spatiotemporal super-resolution microscopy has remained a challenge. Attempts in the past either restrict time resolution^2,3^ or concentration^4,5^, require specialized instrumentation^6,7^, or need specialized sample labeling^8,9^.

While simultaneous multi-parametric fluorescence detection (MFD) has been established for point-measurements^10,11^, such attempts in an imaging mode have been hampered by lack of strategies that can bridge the limitations imposed by spatial and temporal resolution requirements and by the computationally expensive data evaluation procedures required to treat the large data sets. Here, we overcome these problems by acquiring images with high sensitivity and high-speed, using low laser powers at physiological concentrations with genetically encoded labels from the cell-biology fluorophore toolbox^12^, using commercially available cameras and applying GPU-based data processing. We therefore concentrate in this work on the use of standard equipment supported by computational analysis techniques that allow us to simultaneously extract high spatial and temporal resolution from single data sets, in real time.

The datasets are evaluated by a combination of spectroscopy and super-resolution techniques that include: imaging fluorescence correlation spectroscopy (Imaging FCS)^13–15^ to measure dynamics; FCS diffusion law analysis to obtain information about the dynamic sub-resolution molecular organization^16^; Number and Brightness analysis (N&B)^17^ to determine oligomerization or aggregation states; and super-resolution radial fluctuation microscopy (SRRF)^18^ to obtain super-resolved images (Fig. 1).

**Fig. 1:**
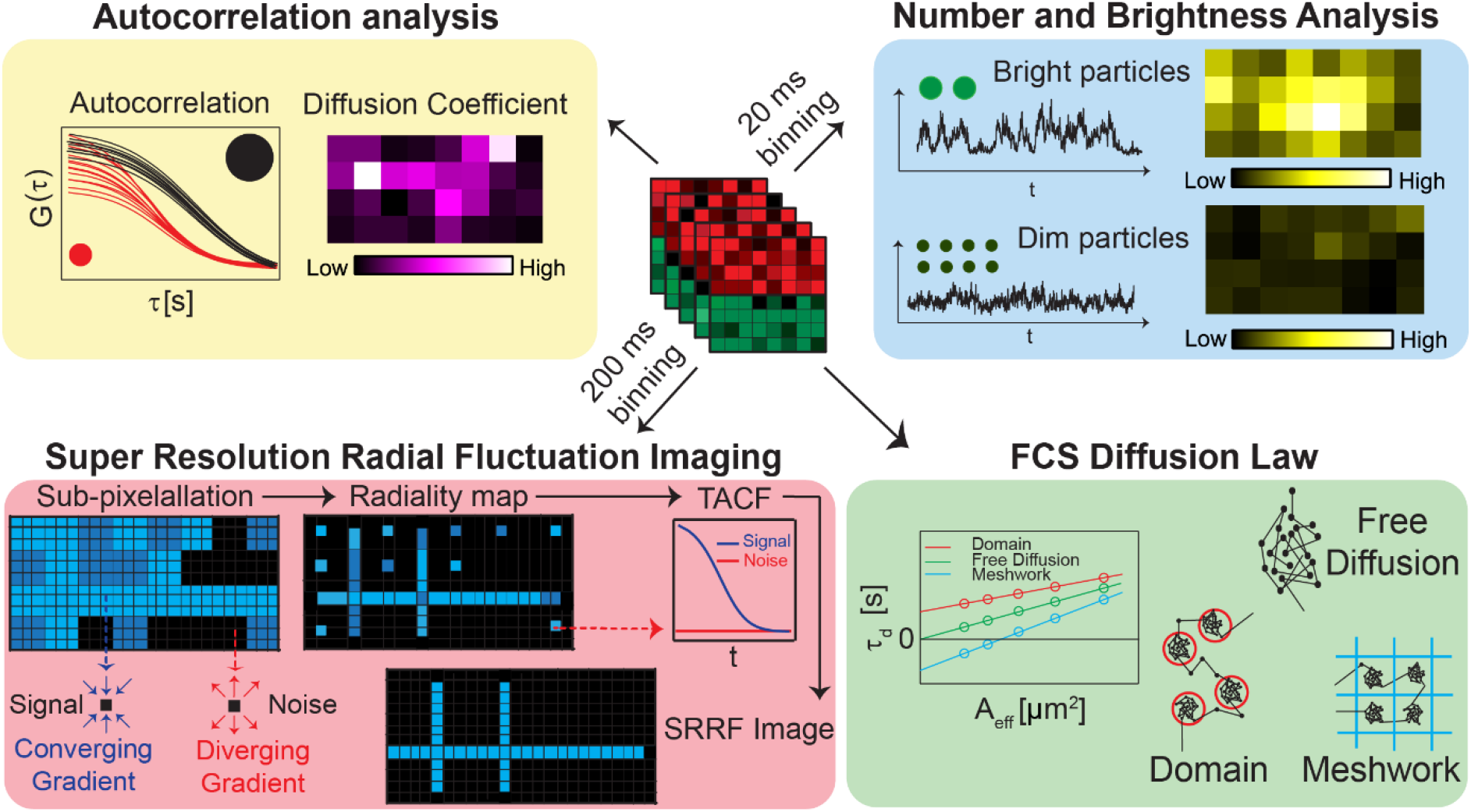
Multi-parametric analysis from a single fluorescence data set: The various analyses performed on a single fluorescence data set are shown here: autocorrelation analysis to determine diffusion coefficient (top left), number and brightness analysis to determine particle brightness (top right), super resolution radial fluctuation imaging to resolve structures (bottom left), and FCS diffusion law to determine protein localization (bottom right).

SRRF^18^, a computational super resolution technique with its roots in super-resolution optical fluctuation imaging (SOFI^19^) yields images resolved beyond the diffraction limit by performing a SOFI analysis on radiality stacks. Imaging fluorescence correlation spectroscopy^13,14^ is a single molecule sensitive ensemble-based method that yields spatially resolved diffusion maps. We performed FCS diffusion law^16^ analysis for the determination of diffusion modes and the sub-resolution organization of the diffusing particles under investigation. In the case of Imaging FCS, the autocorrelation function (ACF) at different lag times is determined from the time-varying intensity at each pixel. In the case of Number and Brightness (N&B)^20^ analysis, only the mean and variance of the time-varying fluorescence intensity function is computed. The concentration and brightness at each pixel are estimated from the computed mean and variance. A comparison of the brightness of a particle with the brightness of a monomer with the knowledge of the probability of a fluorophore to be fluorescent allows estimating the oligomerization state of the particle.

Using a recently published GPU-based algorithm for SRRF^18^ and a newly developed GPU accelerated version of Imaging FCS, N&B and FCS diffusion law analysis (Fig. 1), an ImageJ plugin developed in our group^15^, we achieve measurements with high spatial information (~60 nm) and temporal dynamics (≤ 2 ms) from the exact same data within ~5 min, including measurement time using a commercially available total internal reflection fluorescence (TIRF) microscope. We demonstrate the utility of multi-parametric measurements to monitor the super-resolved structure and dynamics of two different biomolecules, namely LifeAct and epidermal growth factor receptor (EGFR). For this purpose, we recorded image stacks of mApple labeled EGFR (EGFR-mApple) and EGFP labeled LifeAct^21^ on whole cells using EMCCD or sCMOS cameras for detection (50,000 frames at 2 ms time resolution covering areas as large as 128×128 pixels). For the simultaneous acquisition of the mApple and EGFP signals on two halves of one camera, we used a wavelength-based image splitter.

We investigate the localization, super-resolved structure and dynamics of LifeAct, a 17 amino-acid actin binding peptide^21^ demonstrating that spatiotemporal super-resolution can be achieved on one data set measured in one color. Furthermore, we analyzed EGFR receptor dynamics and organization and the cytoskeletal structure on CHO-K1 cell membranes showing that two-color measurements provide additional knowledge that could not be obtained in super-resolution and dynamic measurements separately.

## Results

The acquisition of data at the experimentally best possible temporal and spatial resolution of 50,000 frames at 128×128 pixels at 16 bit per pixel results in files of 1.6 GB size, posing a serious computational challenge for pixel-wise analysis by super-resolution and spectroscopy approaches. We therefore employed a GPU to reduce computational times. A comparison of the time taken for calculating and fitting autocorrelation, diffusion laws and N&B using a CPU and GPU is shown in Fig. S1 for varying sizes of input areas. The achievable improvement is dependent on the total number of pixels being evaluated. Below 160 pixels, GPU processing is slower due to the time required for data transfer to and from the GPU. From about 1,000 pixels onwards we get an improvement of at least a factor 10, depending somewhat on the exact operations. We achieved a maximum acceleration of a factor 38 in the case of N&B analysis of areas above 20,000 pixels (Fig. S1).

Typical measurement times are in the order of 100 s where the sample might show photobleaching. Inclusive of the photobleaching correction, the processing times of ACFs, FCS diffusion law and N&B analysis on a 128×64×50,000 pixel data are 763, 1374, and 99 s with CPU, and 67, 186, and 27 s with GPU evaluation, respectively. The GPU computation times are on the same order as that of the measurement time and can be performed even during acquisition.

Next, we optimized acquisition and evaluation parameters for the various techniques. While results do not depend on the camera, parameters need to be optimized for each camera model as they differ in pixel size, acquisition speed, and achievable SNR.

### Optimization for FCS and FCS diffusion law analysis

The point spread function for the measurements using 488 nm and 561 nm laser was estimated to be 272 nm and 364 nm respectively (Fig. S2). Using these lasers, the measured diffusion coefficients (*D*) of fluorescently labeled molecules diffusing in a supported lipid bilayer are found to be 1.97 ± 0.91 and 2.00 ± 0.34 μm^2^/s (Table S2), similar in range to those reported in the literature^22^.

The FCS diffusion law states that the average transit time through an observation area increases linearly with increase in observation area in the case of a freely diffusing molecule, implying a zero y-intercept in a plot of transition time versus observation area. Non-linearity in the diffusion law plot is typically reported by quantifying the y-intercept of an approximated linear function and is characterized by a non-zero y-intercept. Confined diffusion, for instance, leads to a positive intercept. Here we analyzed the diffusion law from 1×1 to 5×5 (0.48 to 2.10 μm^2^) binning. The diffusion law intercepts (−0.01 s and −0.04 s) were close to zero as expected for a freely diffusing bilayer^23^ (Fig. S2).

### Optimization of N&B parameters

To optimize the brightness parameter, we varied time binning as well as total measurement time to determine the effects of the instrumental parameters on the estimated brightness values. Not all molecules of a fluorescent protein (FP) species are fluorescent due to incomplete maturation, misfolding, photo-bleaching and possible dark states of the fluorophore^24–26^. Hence in order to estimate the oligomerization state of a protein, one needs to estimate the proportion of FPs that are fluorescent. The proportion is estimated by computing the brightness of two different constructs, a monomeric FP and dimeric FP (a single protein consisting of two equal FPs connected by a linker and referred to as a tandem FP). We coupled the first 15 amino-acids of the *RP2* protein to mApple sequences to target it to the plasma membrane^27^, referred to as PMT-mApple (plasma membrane targeted mApple) or PMT-mApple_2_ (plasma membrane targeted mApple-mApple)

While the dimer/monomer brightness ratio (PMT-mApple_2_/PMT-mApple) stabilizes at 40 s total measurement time for all exposure times, it provides consistent values only above 10 ms exposure time (Fig. S3). This is a result of the intensity filtering we use to automatically distinguish between pixels that represent the cell membrane and the background. This distinction improves with exposure time as the difference between cell and background increases with exposure. Here, we used 20 ms exposure time for N&B analysis and 100 s total measurement time for further analysis to maximize accuracy and precision of our results. Using these experimental conditions, we found that the ratio of the brightness of the dimer to that of the monomer (*r*) is 1.55, indicating that 55 ± 1% of mApple are fluorescent (sec. 2.6 in supplement).

### Optimization of SRRF

SRRF was optimized using the actin structure in the green wavelength channel (Fig. S4). Data show that the spatial resolution in SRRF improves with time binning and decreasing pixel size in sample space. At 2 ms time resolution and a pixel size of 120 nm, the full width at half maximum (FWHM) recovered by SRRF for actin structures is 186 ± 9 nm. The FWHM reduces to 59 ± 5 nm with a pixel size of 120 nm and a time binning of 200 ms. Further binning in time does not improve the FWHM but is prone to the creation of artefacts (Sec. 4 in the supplement). The resolution in SRRF images has also been measured using Fourier ring correlation (FRC)^28^ and peak to peak distance (p2p). In our study, the FRC and P2P were found to be 78-145 nm and 120-192 nm (Table S3) respectively for actin fibres.

### Simultaneous SRRF and FCS

Here, we demonstrate that we can study both structure and dynamics of actin on cells from a single data set recorded in a single wavelength channel (Fig. 2). A comparison of the actin structure, provided by TIRF and SRRF images (Fig. 2a-e), and its dynamics, obtained from Imaging FCS (Fig. 2f-h), enables the investigation of structure-dynamics correlations. Evaluations in this section were made on an EMCCD using 200X magnification and 2×2 binning unless stated otherwise.

**Fig. 2:**
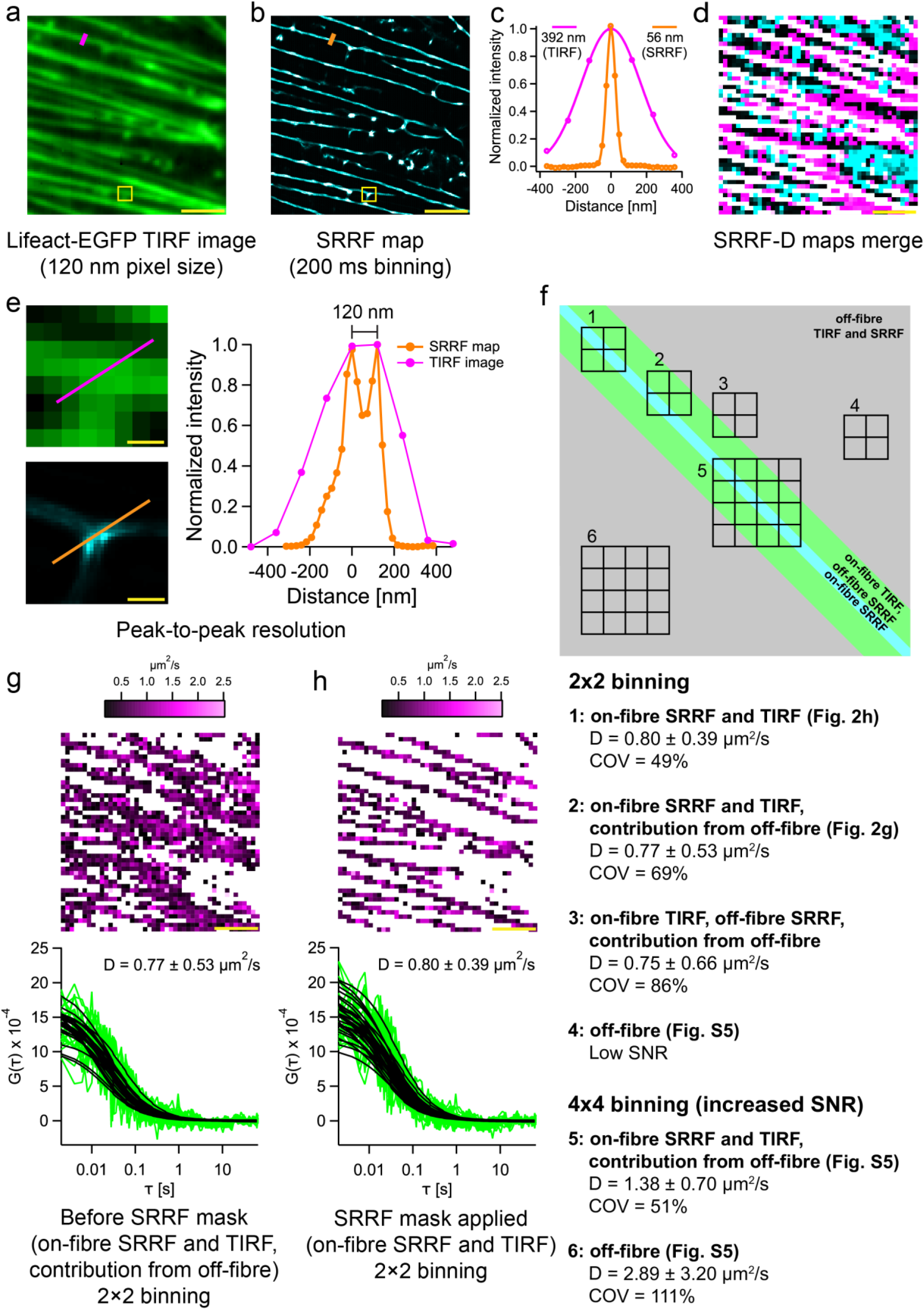
Multi-parametric analysis from a single channel fluorescence data set (CHO-K1 cell labeled with LifeAct-EGFP): **(a)** TIRF image of CHO-K1 cell expressing lifeact-EGFP at 200X magnification. **(b)** SRRF map (200 ms binning; refer Fig. S5) of the cell in (a). **(c)** Normalized intensity profile across an actin fibre before and after SRRF microscopy. Thickness (FWHM of the Gaussian fit) of the actin fibre before SRRF (magenta curve) = 392 nm; measurement point marked with magenta line on image (a). Thickness of the actin fibre after SRRF (orange curve) = 56 nm; measurement point marked with orange line on image (b). **(d)** Merge of the *D* and SRRF maps. The SRRF, *D* and correlated pixels are colored cyan, magenta and white, respectively. **(e)** The enlarged views of the yellow boxes in images (a) and (b) are shown here. The intensity profile below shows the actin fibre branching point (indicated by the orange line on the enlarged SRRF image). The peak-to-peak resolution is 120 nm at this point. The pixel sizes reported are after magnification (refer Table 2). **(f)** Schematic showing the on and off-fibre areas at different bin sizes. **(g)** The *D* map after thresholding (0.2-100 μm^2^/s; *D* = 0.77 ± 0.53 μm^2^/s; refer case 2 in (f)). The coefficient of variation (COV-ratio of standard deviation to the mean) is 69%. Representative ACFs from this map are shown below. (**h)** The *D* map generated after retaining only the pixels that overlap with the SRRF map and removing the remaining pixels shows an average *D* = 0.80 ± 0.39 μm^2^/s (refer case 1 in (f)). The COV is 49%. Representative ACFs from this map are shown below. The scale bars shown in yellow measure 2.5 μm in images (a)-(e) and 250 nm in (f). Mean ± SD is reported here.

The use of SRRF led to an improvement in the resolution of actin fibres. The thickness of actin fibres as quantified by the FWHM of a Gaussian function fitted to the intensity profile was found to be 399 ± 31 nm (Fig. 2a and c, S4) prior to SRRF analysis. SRRF processing led to a 7-fold improvement in the FWHM of actin fibres (59 ± 5 nm, Fig. 2b and c, S4). The resolution as measured by the peak to peak distance was found to be 128 ± 14 nm (Fig. 2e). The diffusion coefficient and SNR of the ACFs vary widely across the image. In general, we find two different cases: a) ACFs in pixels that are along the fibres as judged by the TIRF image (on-fibre SRRF and TIRF, contribution from off-fibre) yield correlations with good SNR and a diffusion coefficients of *D* = 0.77 ± 0.53 μm^2^/s (2f-case 2, Fig. 2g); b) ACFs in between the fibres (off-fibre, Fig. 2f-case 4) exhibited two diffusion components. The faster diffusing component had poor SNR (Fig. S5) at 2×2 binning and could not be fitted. The slow ill-defined component had diffusion coefficients less than 0.1 μm^2^/s. To improve the SNR and obtain more information about the space between the fibres, we repeated the analysis using 4×4 binning (Fig. S5).

The ACFs *in between fibres* using 4×4 binning again exhibited two diffusion components, the fast, and slow ill-defined component. The slow ill-defined component varies widely and randomly (0.01-0.1 μm^2^/s) and is probably an artefact of sample movement and system fluctuations unrelated to the actin diffusion, as can sometimes be seen in FCS at very low amplitudes^29^. We do not further evaluate this slow component.

The fast component (*D* = 2.89 ± 3.20 μm^2^/s) (off-fibre, Fig. 2f-case 6) was isolated by calculating and fitting the ACF only at short lagtime τ between 2 ms and 0.5 s (Fig. S5). This part of the ACF has high a standard deviation, as the time resolution of the camera is only 2 ms and thus does not capture the full ACF for the fast-moving particles. However, the fast component between the pixels indicates rapidly diffusing particles which could be free LifeAct-EGFP or LifeAct-EGFP bound to G-actin. Running the camera at faster frame rates would allow fully capturing faster diffusing particles but at the same time limits the field of view as fast camera read-out is only possible with smaller regions of interest.

The resulting ACFs on the fibres using 4×4 binning (on-fibre SRRF and TIRF, contribution from off-fibre, Fig. 2f-case 5) showed a single component with *D* = 1.38 ± 0.70 μm^2^/s (Fig. S5). The faster diffusion coefficient along the fibres at 4×4 compared to 2×2 binning results from larger contributions from diffusing particles outside the fibre when using the larger 4×4 area for correlation analysis (Fig. 2f-case 5). This indicates that a precise localization of the fibres and a correct attribution of diffusion data to the fibres is required to improve parameter estimations. The diffusion coefficient judged to be along the fibre (0.77 ± 0.53 μm^2^/s, Fig. 2g) by TIRF images has potentially contributions both from the fibre and neighboring areas (Fig. 2f-case 2), as has already been shown by the comparison of FCS results using 2×2 and 4×4 binning.

Hence, SRRF microscopy with its higher resolution and thus better determination of the actual fibre positions can differentiate between those pixels on the FCS diffusion map that are directly on the fibre (on-fibre SRRF and TIRF, Fig. 2f-case1) and those that lie right next to the fibre although they were originally judged to be on the fibre by TIRF and might contain contributions from both on-fibre and off-fibre areas (on-fibre TIRF, off-fibre SRRF, contribution from off-fibre, Fig. 2-case3). Note that these pixels are different from areas between fibres whose diffusion coefficient was found to be 2.89 ± 3.20 μm^2^/s (Fig. 2f-case 6).

The diffusion coefficient on the SRRF-filtered fibre was found to be 0.80 ± 0.39 μm^2^/s (Fig. 2h, Fig. 2f-case 1). Compared to the value obtained without SRRF filtering (0.77 ± 0.53 μm^2^/s, Fig. 2f-case 2) this is an improvement of the error by a factor ~1.4. In the case of pixels neighboring the fibres, as judged by SRRF images, the diffusion coefficient is 0.75 ± 0.66 μm^2^/s (Fig. 2f-case 3).

The larger error of 0.66 μm^2^/s in the estimate of the neighboring pixels is attributed to the lower SNR of the ACFs which are not directly on the fibre. To make the distinction clear we use the ratio of the standard deviation to the mean of the estimate, i.e. the coefficient of variation (COV). The COV improves from 69% (Fig. 2f-case 2) without SRRF masking to 49% (Fig. 2f-case 1), when selecting only pixels directly above the fibres as judged by the SRRF image. In comparison, the COV of neighboring pixels of the SRRF-filtered fibre (Fig. 2f-case 3) was found to be 86%.

A reduction in SNR leads to an increase in the estimate of the number of particles per pixel^30^. The number of particles is lowest on the fibre and increases in the neighboring pixels (Fig. S6). Before filtering, the average number of particles is found to be 912 ± 523, which corresponds to a COV of 57%. (Table S4). After applying the SRRF mask, the average number of particles not only had a 25% reduction but also had a lower COV of 32%, a reduction of a factor 1.7 in COV.

### Two color combined FCS SRRF

The full extent of the information however is gained when analyzing both wavelength channels. To investigate the interrelation between the actin cytoskeleton structure, which is relatively static on the time scale of our measurements, and EGFR mobility and organization, which is highly dynamic, we optimized the signals by spatiotemporal binning for either FCS and FCS diffusion law, N&B (red channel, Figs. 3a-d, Fig. S7) or SRRF (green channel, Figs. 3e-g) analysis. SRRF analysis led to an improvement of the FWHM of the Gaussian fit to the intensity profile across actin fibres from 465 nm to 97 nm. The FWHM is not as low as that from single channel LifeAct experiments (59 ± 5 nm) since a larger pixel size of 240 nm was used in the dual channel measurements.

**Fig. 3:**
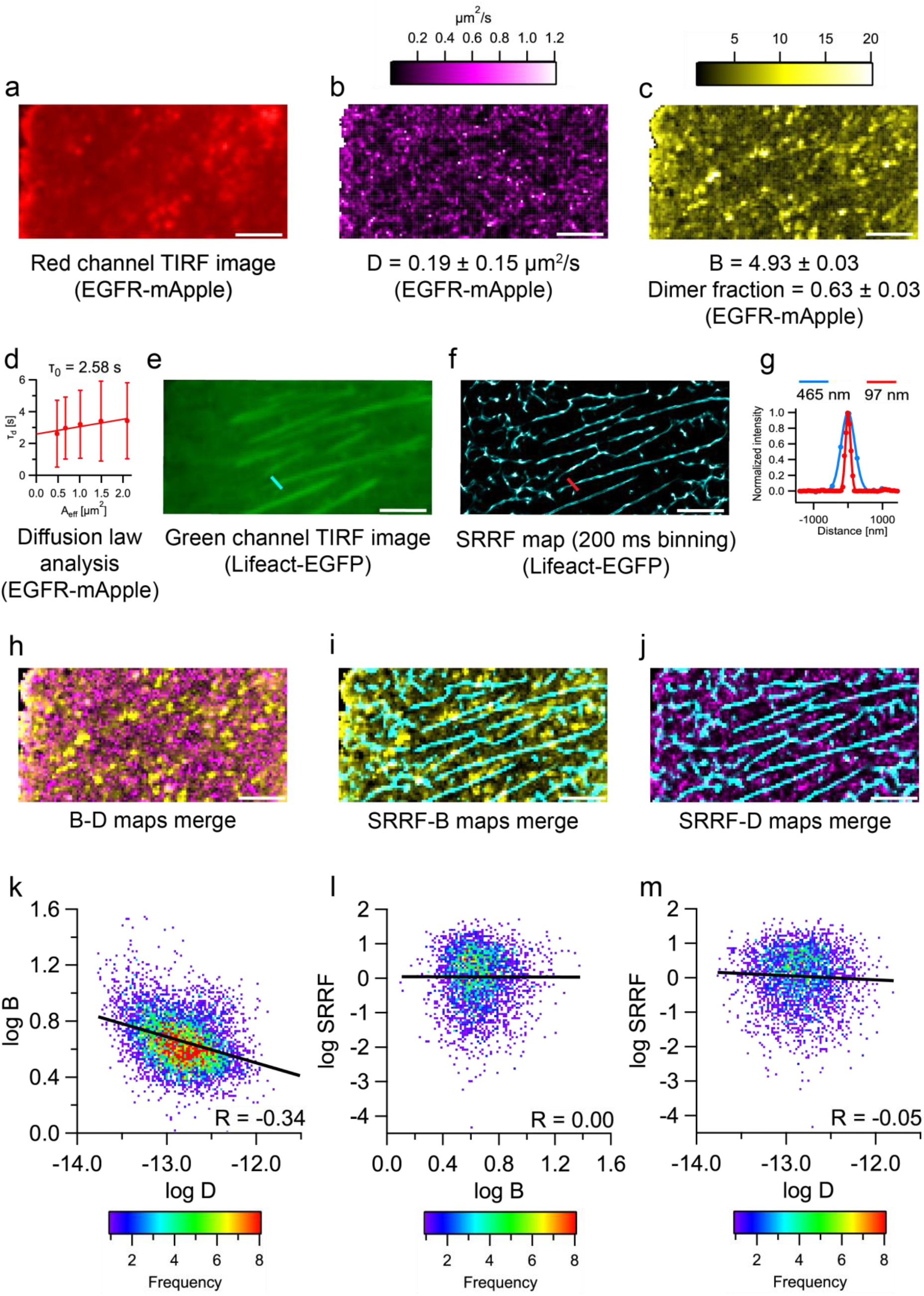
Multi-parametric analysis from a single dual-channel fluorescence data set. **(a)** TIRF image of EGFR-mApple. **(b)** Diffusion map of EGFR-mApple (*D* = 0.19 ± 0.15 μm^2^/s). **(c)** Brightness map of EGFR-mApple (*B* = 4.93 ± 0.03, dimer fraction = 0.63 ± 0.03; mean ± SEM). **(d)** Diffusion law analysis of EGFR-mApple (refer Fig. S10 for more details); intercept = 2.58 s. **(e)** TIRF image of Lifeact-EGFP. **(f)** SRRF map (200 ms binning) of Lifeact-EGFP. **(g)** Normalized intensity profile across an actin fibre before and after SRRF microscopy. Thickness (FWHM of the Gaussian fit) of the actin fibre before SRRF (blue curve) = 465 nm; measurement point marked with blue line on image (e). Thickness of the actin fibre after SRRF (red curve) = 97 nm; measurement point marked with red line on image (f). **(h)** Merge of *B* and *D* maps of EGFR-mApple. The *B* pixels are coloured yellow, and the *D* pixels are coloured magenta. Correlated pixels between the two maps are coloured white. **(i)** Merge of SRRF (Lifeact-EGFP) and *B* (EGFR-mApple) maps. SRRF pixels are coloured cyan, and *B* pixels are coloured yellow. Correlated pixels between the two maps are coloured white. The SRRF map was spatially binned to the same dimensions as the *B* map. **(j)** Merge of SRRF (Lifeact-EGFP) and *D* (EGFR-mApple) maps of EGFR-mApple. SRRF pixels are coloured cyan, and *D* pixels are coloured magenta. Correlated pixels between the two maps are coloured white. The SRRF map was spatially binned to the same dimensions as the *D* map. **(k)** 2D frequency plot of log *B* vs log *D* values from image (h). **(l)** 2D frequency plot of log SRRF vs log *B* values from image (i). **(m)** 2D frequency plot of log SRRF vs log *D* values from image (j). The scale bars in white measure 5 μm in images (a)-(e), (h)-(j). This figure uses cell 1 in Table 1. The reported values are averages obtained from analysis of an entire cell.

The diffusion coefficient was found to be 0.19 ± 0.15 μm^2^/s with a FCS diffusion law intercept of 2.58 s, indicating intermittent trapping of EGFR (Table 1, Fig. 3). N&B analysis of EGFR-mApple showed an intermediate brightness between PMT-mApple and PMT-mApple_2_, indicating that it contains a mixture of EGFR monomers and at least dimers (Table 1, Fig. S7). The average brightness corresponds to an *r* value of 1.35 which translates to 63 ± 3 % dimers. The large error associated with the diffusion coefficient is attributed to the inherent heterogeneity in the cell membrane. The various factors contributing to the heterogeneity are investigated using a multi-parametric analysis.

**Table 1:** Multi-parametric analyses of CHO-K1 cells labeled with LifeAct-EGFP and EGFR-mApple

| Cells | Number of Pixels | $D$<br>[$\mu\text{m}^2/\text{s}$] | $B^*$ | Fraction of monomers in dimer ($m_e$)* | $R$<br>($\log B$,<br>$\log D$) | $R$<br>($\log \text{SRRF}$,<br>$\log B$) | $R$<br>($\log \text{SRRF}$,<br>$\log D$) | FWHM of Gaussian fits to normalized Intensity profiles from SRRF<sup>@</sup><br>[nm] | Diffusion law intercept<br>[s] |
| --- | --- | --- | --- | --- | --- | --- | --- | --- | --- |
| 1 | 6690 | $0.19 \pm 0.15$ | $4.93 \pm 0.03$ | $0.63 \pm 0.03$ | -0.34 | 0.00 | -0.05 | $134 \pm 26.70$ | 2.58 |
| 2 | 3811 | $0.21 \pm 0.19$ | $5.28 \pm 0.03$ | $0.80 \pm 0.03$ | -0.19 | -0.06 | -0.01 | $124.66 \pm 17.01$ | 2.20 |
| 3 | 6720 | $0.31 \pm 0.31$ | $4.97 \pm 0.03$ | $0.65 \pm 0.03$ | -0.40 | 0.02 | -0.08 | $129 \pm 14.76$ | 0.76 |
| 4 | 6061 | $0.11 \pm 0.09$ | $4.78 \pm 0.03$ | $0.56 \pm 0.03$ | -0.50 | -0.07 | 0.00 | $134.66 \pm 8.25$ | 5.16 |
| 5 | 5119 | $0.26 \pm 0.24$ | $4.03 \pm 0.03$ | $0.18 \pm 0.02$ | -0.22 | -0.03 | -0.01 | $141.33 \pm 18.26$ | 1.59 |
| 6 | 3534 | $0.27 \pm 0.25$ | $4.95 \pm 0.06$ | $0.64 \pm 0.03$ | -0.25 | -0.07 | 0.05 | $112 \pm 13.95$ | 1.65 |
| **Mean** |  | **$0.23 \pm 0.22$** | **$4.82 \pm 0.04^*$** | **$0.58 \pm 0.03^*$** | **$-0.32 \pm 0.11$** | **$-0.03 \pm 0.03$** | **$-0.02 \pm 0.04$** | **$129.28 \pm 19.72$** | **$2.32 \pm 1.52$** |

\*The reported values are mean ± SEM. In all other cases, the values are mean ± SD,<sup>@</sup> The reported values are mean ± SD from three positions in each cell measurement.

EGFR-mApple expressing cells show both regions of uniform intensity and regions exhibiting visible clustering (Fig. 3a). Analysis of the regions of homogeneous brightness in the mApple channel by Imaging FCS and the FCS diffusion law resulted in a diffusion coefficient of *D* = 0.24 ± 0.16 μm^2^/s and a positive diffusion law intercept of 1.81 s (Fig. S8), indicating transiently trapped diffusion and possibly EGFR oligomerization or clustering beyond the resolution of our system^31^. The existence of oligomers is corroborated by N&B analysis of the same regions. EGFR-mApple showed a brightness intermediate between monomers and dimers (Fig. S7), suggesting that 18 ± 6% of receptors are found in dimers, although we cannot exclude the existence of small amounts of higher oligomers^32^.

When analyzing the regions with visible clusters present, we obtain a *D* = 0.11 ± 0.08 μm^2^/s (Fig. S8) via FCS and a *r* = 1.6 via N&B analysis. In this case, a simple dimer model is not reasonable anymore. Given that only 55% of mApple molecules are fluorescent based on calibration experiments, *r* > 1.55 indicates the presence of oligomers larger than dimers (Sec. 2.6 in the supplement). For instance, the value of *r* = 1.6 would translate into 71% monomers and 29% trimers, or 94% dimers and 6% trimers. It is important to note that if more than two species are present, extra experimentally measured brightness values are necessary. Nevertheless, the equations provide information on the existence of higher oligomers and determine limits on their fractions.

To investigate the membrane organization, we used diffusion law analysis over different areas of CHO-K1 cells expressing EGFR-mApple. There were four kinds of areas which include the presence or absence of cytoskeleton or clusters. The positive intercepts for all entries in Table S5 indicated that EGFR is localized in lipid domains independent of the presence or absence of cytoskeleton or clusters (Fig. S8, Table S5). Furthermore, there was no statistically significant difference in intercepts obtained either in the case of regions with or without clusters or in the case of regions with or without actin (Table S5, p > 0.14). The diffusion law intercept of the entire cell is similar to that of the region containing both actin and cytoskeleton.

Since all evaluations stem from the exact same data set we overlap the parameter maps and pairwise analyze their scatter plots to determine the interdependence of protein location with its dynamics. We performed a logarithmic transformation of the entire dataset to identify power law scaling between parameters (Fig. 3h-m).

Using whole cell data, we find a negative correlation between particle brightness and diffusion coefficient (Pearson’s correlation coefficient (*R*) = −0.34, Table 1 and Fig. 3h, k), indicating that brighter particles diffuse slower than dimmer particles. This is unexpected since membrane diffusion is only weakly dependent on size^33^. This implies either that EGFR oligomers are located in a more viscous membrane environment, e.g. lipid domains^34^ or clathrin-coated pits, or that we detect multiple EGFR molecules in larger complexes^32,35,^. We also obtain a negative correlation (*R*= −0.83) between diffusion coefficient and the positive diffusion law intercept indicating that trapping is responsible for lower EGFR mobility as expected (Table 1).

Brightness and SRRF do not show any correlation (*R* = 0.00, Fig. 3 i,l), implying that clustering is not directly linked to the cytoskeleton. Similarly, we do not see any correlation between EGFR diffusion and cytoskeletal structures (*R* = −0.05, Fig. 3 j,m), unlike in the case of LifeAct. This also verifies our earlier observation that diffusion law intercepts are independent of the presence or absence of actin. Similar results were obtained by performing the experiment using a sCMOS camera as detector (Figs. S9-S11).

## Discussion

Our approach allows the simultaneous determination of super-resolved structure and millisecond molecular dynamics from the exact same pixels over a whole image without any special sample preparation. This provides a new tool to determine correlations between molecular structure and dynamics using a single or better two colors as illustrated by LifeAct and EGFR. It is important to note that, although here we have used a combination of SRRF, simultaneous multiparametric spatiotemporal microscopy is also possible with other computational super resolution tools, e.g. 3B analysis^36^.

Using the same dataset, we performed combined Imaging FCS and SRRF measurements. In our study, the FRC and P2P were found to be 78-145 nm and 120-192 nm respectively for actin fibres similar to previously published resolutions obtained with SRRF, which obtained an FRC of 108 nm^37^, and a P2P distance of 64 nm for actin in fixed cells^18^.

The use of simultaneous SRRF and Imaging FCS measurements led to an improvement in the precision of the estimate of the mobility and number of particles of LifeAct-EGFP molecules when compared to estimates obtained only from Imaging FCS. Initially, the gain in accuracy and precision using SRRF filtering to determine localized diffusion coefficients seems to be small. However, any gain of accuracy and precision within live cells is important as it provides better values and limits for modelling approaches. But even more importantly, the correlation between fibre position as judged by SRRF and SNR and the COV of FCS provide mutual support for the two techniques. The SRRF position is corroborated by the lower COV of FCS, and vice versa. This allows a clearer distinction between diffusion on and off the fibres. The slower diffusion coefficient on the fibres implies binding and unbinding of LifeAct and thus a slowdown in mobility, while the faster diffusion coefficient of lifeact between the fibres indicates free diffusion of LifeAct or LifeAct bound to globular actin^21^.

In the case of two-color measurements, in order to estimate the oligomerization of EGFR, we first estimated the proportion of fluorescent molecules of mApple. We found that 55 ± 1% of mApple are fluorescent which is well within the upper range of literature values for red FPs of 20-70%^25^. Using this calibration, we find that 63 ± 3 % of transfected EGFR molecules exist as dimers in CHO-K1 cells, which do not have endogenous expression of EGFR^38^. The existence of preformed oligomers is consistent with literature with up to ~65% dimerization previously detected^32,38,39^ in the same cell line.

FCS diffusion law analysis showed that EGFR has a positive intercept indicating that it is partitioned into domains. We observed brighter particles diffuse slower compared to dimmer particles whereas there was no correlation between the diffusion of EGFR and the underlying cytoskeleton. This indicates that the domain partitioning of EGFR is primarily determined by EGFR-membrane interactions, possibly sub-resolution clustering and inter-EGFR interactions but independent of EGFR-actin interactions. As it has been shown previously that EGFR dynamics changes with cytoskeleton disruption^40^, our observations suggest that it is not the static cytoskeleton itself that changes EGFR dynamics but that the influence is indirect and cytoskeleton coupling to the plasma membrane and resulting membrane changes are responsible for changes in EGFR dynamics^41^. However, we cannot exclude the possibility that our spatial resolution for the dynamics, even with SRRF filtering, is not sufficient to detect more subtle EGFR/actin interactions and this issue deserves closer inspection in the future.

sCMOS cameras can be used as alternative detectors to EMCCDs although there are some differences. EMCCD cameras have somewhat better SNR at low light levels and are thus better detectors under these circumstances. However, they have also much larger pixels which is counterproductive when spatial super-resolution is to be achieved, unless corrected by changing the magnification. sCMOS cameras on the other hand can be read-out at least one order of magnitude faster than EMCCDs and thus can provide access to faster diffusing particles, but in that case need higher laser power to obtain sufficient signal^42,43^. Despite these differences, both detectors provide the same absolute parameter values in our measurements albeit with different SNR (EMCCD-6.1, sCMOS-1.4).

## Conclusion

Living systems are highly dynamic with processes happening on spatial scales well below the optical diffraction limit and on time scales on the millisecond scale or faster. Therefore, the extraction of the maximum information available on biological processes requires the simultaneous acquisition of data with high spatiotemporal resolution. This poses particular problems to data recording and data evaluation strategies that allow data treatment in an acceptable time frame, ideally in real-time. We used GPU-based data evaluation and modern camera technology to optimize data evaluation of fluorescence microscopy images by selected spatial and temporal binning to extract multiple physicochemical parameters in parallel from a single data set, almost in real time, using standard and widely available instrumentation. Parameter correlations and the lack thereof provide information not available on sequentially acquired data due to the dynamical nature of biological samples as shown on the example of EGFR. This approach is easily extendable to other fluorescence parameters, does not require specialized instrumentation, and thus is immediately applicable to a wide range of situations. We have shown that on a TIRF microscope but the same strategy is applicable to any illumination system that can efficiently capture a cross section of a 3D sample. Especially the lack of requirement of any customized instrumentation and the provision of the necessary evaluation software makes this approach immediately applicable to a wide set of researchers.

## Methods

### Sample preparation

CHO-K1 (Chinese Hamster Ovary) cells (CCL-61) were obtained from ATCC (Manassas, Virginia, USA) and were cultured in Dulbecco’s Modified Eagle Medium (DMEM/High glucose with L-glutamine, without sodium pyruvate – SH30022.FS; HyClone, GE Healthcare Life Sciences, Utah, USA) supplemented with 1% penicillin-streptomycin (Gibco, Thermo Fisher Scientific, Massachusetts, USA), and 10% fetal bovine serum (FBS; Gibco, Thermo Fisher Scientific, Massachusetts, USA) at 37 °C in a 5% (v/v) CO_2_ environment (Forma Steri-Cycle CO_2_ incubator, Thermo Fisher Scientific, Massachusetts, USA).

The construction of the plasmids PMT-mApple, PMT-mApple_2_ and EGFR-mApple are detailed in the supplement. Lifeact-EGFP plasmid was a gift from Prof. Wu Min (NUS, Singapore). All the plasmids were amplified using ZymoPURE II Plasmid Midiprep Kit (D4201; Zymo Research, California, USA) and their concentration and purity were confirmed by a UV-Vis spectrophotometer (NanoDrop 2000, Thermo Fisher Scientific, Massachusetts, USA). The plasmid amounts used for transfection were 100 ng for PMT-mApple, PMT-mApple_2_ and Lifeact-EGFP, and 1 μg for EGFR-mApple.

For transfection, cell cultures that were ~90% confluent were used. The spent media removed from the culture flask was discarded. The flask was washed twice with 5 ml PBS (phosphate buffered saline; without Ca^2+^ and Mg^2+^). 2 ml Trypsin-EDTA (0.25%; Gibco, Thermo Fisher Scientific, Massachusetts, USA) was added and the flask was incubated at 37°C for 2-3 minutes to detach the cells. 5 ml culture media was added to the flask to inhibit trypsin. The media containing the detached cells was collected in a falcon tube and centrifuged at 1,000 rpm for 3 minutes. The supernatant was discarded and the cell pellet was resuspended in 5 ml PBS. Cell counting was done using a cell counter (Bio-Rad, Singapore). The required number of cells was aliquoted into falcon tubes and centrifuged at 1,000 rpm for 3 minutes. The supernatant was discarded and the cells were resuspended in R buffer (Neon Transfection Kit, Thermo Fisher Scientific, Massachusetts, USA). Suitable amounts of Lifeact-EGFP and EGFR-mApple (or PMT-mApple, or PMT-mApple_2_) (100 ng for Lifeact-EGFP; 1 μg for EGFR-mApple) plasmids were mixed with the cells for co-transfection. The cells were electroporated using Neon Transfection system (Thermo Fisher Scientific, Massachusetts, USA) according to the manufacturer’s protocol (electroporation settings: pulse voltage = 1,000 V, pulse width = 30 ms, and pulse no. = 2). After transfection, the cells were seeded onto culture dishes (MatTek, Massachusetts, USA) containing DMEM (supplemented with FBS). The cells were incubated at 37°C and 5% CO_2_ for 36-48 hours before measurements.

Before EGFR measurements, the cells were washed with HBSS (Hank’s Balanced Salt Solution, with Ca^2+^ and Mg^2+^; #14025134; Gibco, Thermo Fisher Scientific, Massachusetts, USA) and starved in DMEM not containing phenol red (#21063029; Gibco, Thermo Fisher Scientific, Massachusetts, USA) for at least 4 hours. To avoid internalization of EGFR, internalization inhibitors were added to the cells 30 minutes before the measurements. The internalization inhibitors used were 2 mM NaF, 10 mM NaN_3_ and 5 mM 2-deoxy-D-glucose (Sigma-Aldrich, Singapore).

### Instrumentation

The TIRF microscopy set-up included an inverted epi-fluorescence microscope (IX83, Olympus, Singapore), a motorized TIRF illumination combiner (IX3-MITICO, Olympus, Singapore), and a dual-emission image splitter (OptoSplit II; Cairn Research, Faversham, UK). We used either an electron multiplying charge-coupled device (EMCCD; iXon^EM^+ 860, 128×128 pixels, Andor, Oxford Instruments, UK) camera or a scientific complementary metal oxide semiconductor (sCMOS; Sona 4.2B-11, Andor, Oxford Instruments, UK) for detection. 488 nm (LAS/488/100, Olympus, Singapore) and 561 nm (LAS/561/100, Olympus, Singapore) lasers were connected to the TIRF illumination combiner. We used a 100X, NA 1.49 oil-immersion objective (Apo N, Olympus) and a magnification changer slider (IX3-MITICO, Olympus, Singapore) to increase magnification two-fold to 200X where required. For the cell measurements, 37°C temperature and 5% CO_2_ atmosphere were maintained using an on-stage incubator (Chamlide TC, Live Cell Instrument). The laser power used was 100 μW for the 488 nm laser and 900 μW (EMCCD) for the 561 nm laser (as measured at the back aperture of the objective).

For the dual-channel measurements, the fluorescence light was passed through a dichroic (Di01-R488/561; Semrock) and split by the image splitter on two halves of the camera chip. The image splitter was fitted with an emission dichroic (FF560-FDi01; Semrock) and band-pass filters (510AF23 and 685ALP, respectively; Omega Optical). A bright-field image of a stage micrometer was used to align the image splitter. This was done in μManager (https://micro-manager.org). The image was aligned in both the channels following the manual instructions. To check how good the alignment was, a self-written program in μManager was used to find the similarity in both channels. We considered the channels sufficiently aligned if the similarity was ≥ 95%.In the case of EMCCD, the measurements were done by recording a stack of 50,000 frames of 128×128 pixels at 500 frames per second (fps) (for cell measurements)/1,000 fps (for bilayer measurements). Andor Solis was used for image acquisition. The kinetic mode of image acquisition was used and the ‘baseline clamp’ was always used to minimize the baseline fluctuation. The camera was operated using 10 MHz pixel readout speed. Maximum analog-to-digital gain was set to 4.7 and 0.45 μs vertical shift speed were used. The EM gain used was 300.

### Data analyses

The data analyses was performed on a computer with the following configuration – Windows 10 Home 64-bit operating system, Intel® Core™ i7-7800X CPU @ 3.50 GHz processor, 32 GB RAM, NVIDIA Titan X_p_ GPU with 3840 CUDA cores and 12.3 GB memory.

#### FCS

The image stacks from the cell measurements were loaded in Imaging FCS^44^ 1.52 plugin for Fiji and the ACFs calculated. The values used for the parameters were: frame time = 0.002 s, correlator (p, q) = (16, 12), pixel size = 24 μm, NA = 1.49, λ_1_ = 565 nm (red channel). The EMCCD data analysis was performed at 1×1 binning for 100X magnification. Bleach correction^45^ was performed with a polynomial of order 8. ACFs were fitted with a one-component diffusion model^46^.

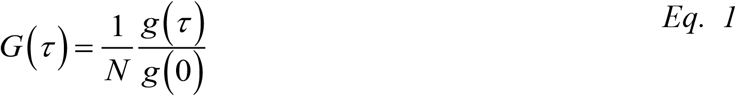

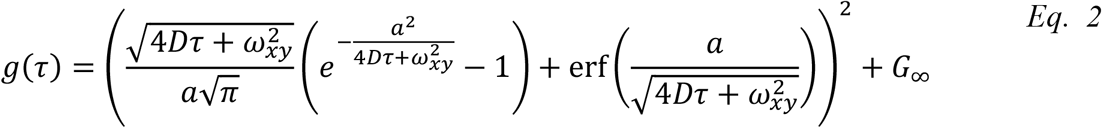

where *a* is the pixel size, *τ* is the lag time, *N* is the number of particles, *D* is the diffusion coefficient, *ω_xy_* is the PSF (xy) (*1/e^2^* radius) direction. To exclude outliers or non-converged fits, only data with 0.01 < *D* < 10 μm^2^/s were considered for further analysis. The point spread function was estimated using the method described here^47^.

#### FCS diffusion law

The image stack was analysed using the Imaging FCS 1.52 plugin for Fiji. The values used for the parameters were: frame time = 0.002 s, correlator (p, q) = (16, 12), pixel size = 24 μm (EMCCD), magnification = 100, NA = 1.49, λ1 = 565 nm (red channel), PSF (xy) = 0.96 (EMCCD red channel) and bleach correction = polynomial of order 8. In the “Diff. Law” tab, the diffusion law plot was generated and fit with a straight line for square binning 1 to 5 (EMCCD).

#### N&B

The 2 ms, 50,000 frames image stack was temporally binned (sum binning of 10 frames each) to a 20 ms, 5,000 frames stack in Fiji. The images were analysed using the Imaging FCS 1.52 plugin. The values used for the parameters were: frame time = 0.02 s, binning = 1×1 (EMCCD), and bleach correction = polynomial of order 8. An intensity filter with a suitable range was set, as defined by the background intensity in an image, to exclude the background pixels and include only the pixels containing the cell. A dark image (image taken with the camera shutter closed) with the same spatiotemporal dimensions as the measurement image was loaded for background correction in the “Bgr NUM” tab. Then in the “N&B” tab, “G1” was selected in the “NB mode” and the “N&B” button in “N&B analysis” was pressed to generate the N&B maps. The N&B equations are defined as^17^:

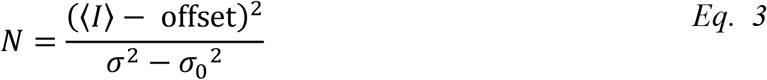

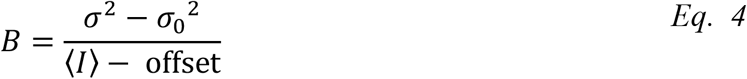

where *N* is the apparent number of particles in the observation volume, ⟨*I*⟩ is the average intensity, offset is the intensity offset of EMCCD, *σ*^*2*^ is the variance of the signal, *B* is the apparent brightness of a particle, and *σ*_*0*_^*2*^ is the variance of the readout noise in the EMCCD. The offset and *σ*_*0*_^*2*^ can be obtained from a dark image captured by the EMCCD. The variance is typically affected by shot noise, and hence we used the covariance. This is referred to as G1 analysis.

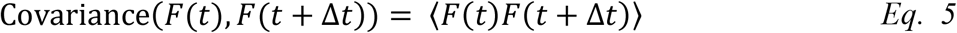

where *F(t)* is the fluorescence signal at time *t* and *F(t+Δt)* is the fluorescence signal at time *t+Δt.*

The dimer fraction is determined using the equation below:

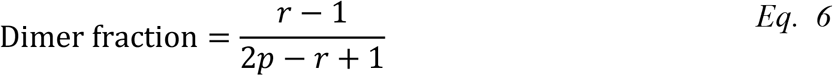

where *r* is the ratio of the brightness of EGFR to the brightness of monomer, *p* is the proportion of the molecules which are fluorescent. The procedure to determine *p* based on the brightness of monomer and dimer is provided in supplement. The error propagation to estimate the error associated with the dimer fraction is also provided in supplement.

#### SRRF

The 2 ms, 50,000 frames image stack was temporally binned (average binning of 100 frames each) to a 200 ms, 500 frames stack (for Fig. S4 other temporal binnings were also used) in Fiji using the Image → Transform → Bin command sequence. The NanoJ-SRRF^18^ plugin was used. The “SRRF analysis” option in the plugin was chosen. The default settings were used, with an addition – the “Do Gradient Smoothing” (“Show Advanced Settings” → “Radiality” → “Do Gradient Smoothing”) was also activated. To determine the fibre thickness, the straight line tool in Fiji was used to draw a line segment across a fibre in the generated SRRF maps. The “Plot Profile” function was used to generate an intensity histogram, which was fitted using the “Curve Fitting” tool in Fiji. The curve fitting function used was a Gaussian given by the equation below.

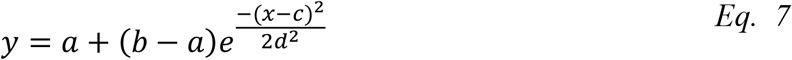

where *a* is the offset, *b-a* is the height at the centre *c,* and *d* is the standard deviation of the Gaussian. The SRRF maps had black borders so the image was cropped to get rid of them. As a result, the original acquired image was cropped before performing FCS, diffusion law and N&B analyses to maintain the same cell area in all the maps (for the image dimensions, refer table 2).

**Table 2:** Acquisition parameters for the various experimental configurations

| Camera | Analysis | Laser wavelength [nm] | Time per frame [ms]* | No. of frames | Image pixel size [nm] | Map dimensions [Pixels] |
| --- | --- | --- | --- | --- | --- | --- |
| EMCCD | FCS | 488 | 2 | 50,000 | 240 | 46x46 |
| **Fig. 2** | SRRF | 488 | 200 | 500 | 24 | 460x460 |
| EMCCD | FCS | 561 | 2 | 50,000 | 240 | 120×56 |
| **Fig. 3** | Diffusion law | 561 | 2 | 50,000 | 240 | 114×56 |
|  | N&B | 561 | 20 | 5,000 | 240 | 120×56 |
|  | SRRF | 488 | 200 | 500 | 48 | 600×280 |

\*: acquisition time was 2 ms; longer times are reached by time binning of frames.

### SNR calculation

For the SNR calculation on the EMCCD and sCMOS, an area of 20×20 pixels was chosen inside and outside a cell. The area outside the cell was used to estimate the background for SNR calculation using the equation below.

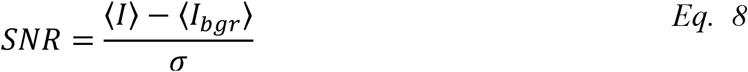

where ⟨*I*⟩ is the average intensity inside the cell (signal), ⟨*I_bgr_*⟩ is the average intensity outside the cell (background), and *σ* is the standard deviation of the signal inside the cell.

### Statistical analysis

The frequency plots in Fig. 3 were generated using the “*Bivariate histogram*” command in IgorPro© (Wavemetrics Inc, Oregon, USA). A linear fit was performed to estimate the correlation coefficient (*R*). Graphpad Quickcalcs was used to perform t-tests. Only those pixels were included in the scatter plots that had valid values for both parameters.

## Code availability

The Imaging FCS 1.52. ImageJ plugin is available at http://www.dbs.nus.edu.sg/lab/BFL/imfcs_image_j_plugin.html or alternatively is included in the ImageJ update site.

## Data availability

The datasets generated during the current study are available from the corresponding author on reasonable request.

## Competing Interests

The authors declare that there are no competing interests.

## Author Contributions

T.W conceived and designed the study. J.S and W.H.T wrote programs for GPU analysis. H.B and X.W.N performed the experiments. J.S and H.B analyzed the data. J.S, H.B, W.H.T and T.W wrote the manuscript. A.R and T.W supervised the study.

