## Supplementary Information for "SRRF ‘n’ TIRF - FCS: Simultaneous spatiotemporal super-resolution microscopy"

### 1. Introduction

There exists a wide variety of fluorescence spectroscopy and microscopy techniques that are able to obtain either spatial super-resolution or high temporal resolution. Unfortunately, spatial super-resolution techniques often require special instrumentation that not only limits their availability but also restricts the possibility to obtain good temporal resolution.

We demonstrate a strategy to perform simultaneous multi-parametric analysis using fast and sensitive cameras (EMCCD and sCMOS) and a combination of super-resolution and spectroscopy techniques – super resolution radial fluctuation (SRRF) imaging to resolve structures, fluorescence correlation spectroscopy (FCS) analysis to determine diffusion coefficient, number and brightness analysis to determine particle brightness, and FCS diffusion law analysis to determine membrane organization (Fig. 1). The system is implemented on a total internal reflection fluorescence (TIRF) microscope.

SRRF[1,2], a computational super resolution technique with its roots in super-resolution optical fluctuation imaging (SOFI[3]) and yields images resolved beyond the diffraction limit by performing a SOFI analysis on radiality stacks. Imaging fluorescence correlation spectroscopy[4,5] is a single molecule sensitive ensemble-based method that yields spatially resolved diffusion maps. A statistical analysis of fluctuations in fluorescence in each pixel of an array detector provides the autocorrelation function in every pixel. Fitting the autocorrelation curve in each pixel to theoretical models yields the diffusion coefficient ($D$) and the number of particles in that pixel ($N$).

We performed FCS diffusion law analysis for the determination of diffusion modes and the sub-resolution organization of the diffusing particles under investigation. For this purpose, the estimated diffusion coefficient at various observation areas are transformed to yield the average transit time through the area. The FCS diffusion law[6] states that the average transit time through an observation area increases linearly with increase in observation area in the case of a freely diffusing molecule. Non-linearity in the diffusion law is typically reported by quantifying the y-intercept of an approximated linear function. A positive intercept is obtained in the case of confined diffusion.

In the case of Imaging FCS, the autocorrelation function at different lag times is determined from the time-varying intensity at each pixel. In the case of Number and Brightness (N&B)[7] analysis, only the mean and variance of the time-varying fluorescence intensity function is computed. The concentration and brightness at each pixel are estimated from the computed mean and variance. To obtain proper estimations of brightness values in N&B analysis, it is important to take correctly account of the background and its variance as produced by the detector, in this case EMCCD or sCMOS cameras as described here[8]. A comparison of the brightness of a particle with the brightness of a monomer with the knowledge of the probability of a fluorophore to be fluorescent enables one to estimate the oligomerization state of the particle.

As a proof of principle, we perform a two-color experiment with EGFR-mApple and LifeAct-EGFP in live CHO-K1 cells. The results demonstrate that we can obtain super-resolution in space (<60 nm) with simultaneously high resolution in time (2 ms) from the exact same data. This allows us to mutually correlate the various parameters and obtain new information on EGFR organization and dynamics in relation to the cytoskeleton.

### 1 Supplementary methods

#### 1.1 Plasmid construction

The N-terminal of the mApple sequence was coupled with the first 45 nucleotides of the *RP2* gene to target it to the plasma membrane[9]. The plasmid containing the pRK-5 vector backbone with a pUC origin of replication, cytomegalovirus (CMV) promoter, SV40 polyadenylation signal site, ampicillin resistance gene cloned with the membrane targeted mApple was obtained from VectorBuilder Inc (Illinois, USA). We refer to this plasmid as plasma membrane targeted mApple (PMT-mApple). To create a double labeled PMT-$mApple_2$ the PMT-mApple plasmid was digested with NheI and HindIII to create the backbone. The plasmid was also separately digested with SpeI and HindIII to get the mApple sequence insert. The backbone and the insert were ligated (this was possible because SpeI and NheI are isocaudomers) using T4 DNA ligase (M0202S; New England BioLabs, Massachusetts, United States) to create PMT-$mApple_2$. The EGFR-mApple plasmid was created by replacing the PMT sequence with EGFR sequence using restriction digestion with AgeI and SpeI and ligation using T4 DNA ligase. The plasmid maps created using SnapGene® software (GSL Biotech LLC, IL, USA) are shown in Fig. S12.

#### 1.2 Supported lipid bilayer (SLB) preparation

All glassware (slides, coplin jars and round-bottom flasks) were cleaned thoroughly using a 10X diluted alkaline cleaning solution (Hellmanex III, Hellma Analytics, Singapore). They were then submerged in the cleaning solution and sonicated (Elmasonic S30H, Elma Schmidbauer GmbH, Singen, Germany) for 30 minutes. This was followed by washing with ultrapure water (Milli-Q, Merck, New Jersey, United States) and another round of sonication for 30 minutes after submerging in 2M $H_2SO_4$. The glassware were then washed and submerged in ultrapure water for a third round of sonication for 30 minutes. After air-drying, the glassware were used for SLB preparation. An O-ring mould was filled with a silicone elastomer (SYLGARD 184 Silicone Elastomer Kit, Dow, Michigan, USA) and cured at 65°C overnight. The O-rings were then removed carefully using forceps, attached to a slide using the silicone elastomer and cured for 3 hours at 65°C. Unused glassware and O-rings were stored in 100% ethanol for later use.

0.5 mM DOPC (1,2-dioleoyl-sn-glycero-3-phosphocholine; Avanti Polar Lipids, Alabama, USA) and 50 nM Rhodamine PE (14:0 Liss Rhod PE, Avanti Polar Lipids, Alabama, USA)/100 nM Lipilight 488 (Idylle, Paris, France) were mixed in a round-bottom flask. The solution was evaporated in a rotary evaporator (Rotavapor R-210, Büchi, Flawil, Switzerland) and a thin lipid film was left behind. The lipid film was dissolved in 2 ml of a buffer solution (10 mM HEPES, 150 mM NaCl, pH 7.4) and sonicated until the solution became clear. 200 µl of the lipid solution was pipetted into an O-ring attached to a slide. The slide was incubated at 65°C for 1 hour to allow the SLB to form by vesicle fusion. Then it was cooled at room temperature (25°C) for 30 minutes. The SLB was washed multiple times with the buffer solution (by removing 100 µl of lipid solution inside the O-ring and adding 100 µl of buffer) to get rid of excess, unfused vesicles and then used for measurements.

#### 1.3 Instrumentation

The TIRF microscopy set up is described in the main text. Apart from the EMCCD described in the main text, here we used also an sCMOS (Sona 4.2B-11, Andor, Oxford Instruments, UK) camera for measurements. The following settings were used for recording the image stacks. The laser power was set to 100 µW for the 488 laser and 19 mW for the 561 nm laser. We recorded stacks of 50,000 frames of 150×300 pixels at 500 fps (for cell measurements) or 1,000 fps (for bilayer measurements). The software Andor Solis was used for image acquisition. The acquisition mode used was "kinetic series". The pixel readout rate was 200 MHz in an overlap readout mode. The rolling electronic shuttering mode and 12-bit dynamic range was used. The acquisition settings for the various experimental configurations are summarized in Table S1.

The point spread function (PSF) calibration measurements were performed with 500 µW laser power at 488 nm laser and 1 mW at 561 nm, as described in the next section.

**Table S1: Acquisition parameters for the various experimental configurations**

| **Camera** | **Analysis** | **Laser [nm]** | **Time per frame [ms]** | **No. of frames** | **Image pixel size [nm]** | **Map dimensions [Pixels]** |
|---|---|---|---|---|---|---|
| **sCMOS** | FCS | 561 | 2 | 50,000 | 220 | 71x71 |
| **Fig. S9** | Diffusion law | 561 | 2 | 50,000 | 330 | 10x10 |
| | N&B | 561 | 20 | 5,000 | 220 | 71x71 |
| | SRRF | 488 | 200 | 500 | 22 | 710x710 |

#### 1.4 Calibration of PSF for Imaging FCS

The instrumental parameters in the fitting model in Imaging FCS are the pixel size and the point spread function (PSF). The PSF is determined in Imaging FCS based on the fact that the estimated diffusion coefficient is independent of the observation area used, as determined by pixel binning, since the diffusion coefficient is an intrinsic molecular property. The diffusion coefficient at various bin sizes are determined for different values of the PSF. The PSF which yields a diffusion coefficient independent of the bin area is the PSF of the system. For the purpose of the calibration we define the PSF as

$$PSF = \frac{\omega_{xy}\lambda}{NA} \qquad \textit{Eq. 1}$$

where $\omega_{xy}$ is the dimensionless scaling factor, $\lambda$ is the wavelength of emission, and $NA$ is the numerical aperture of the detection objective. To calibrate the PSF for the different experimental setups, the relevant bilayer (Rhodamine PE for 561 nm and Lipilight 488 for 488 nm) measurement file of 20×20 pixels was loaded into the program. The values used for the parameters were: frame time = 0.001 s, correlator ($p$, $q$) = (16, 9), pixel size = 24 µm (EMCCD)/11 µm (sCMOS), $NA$ = 1.49, $\lambda$ = 506 nm (green channel)/565 nm (red channel). The program computed and displayed a plot of the $D$ values for various combinations of PSF and binning values.

#### 1.5 Data preprocessing

In the case of 200X magnification, we used 2×2 binning for Imaging FCS analysis on the EMCCD to maintain a good signal-to-noise ratio (SNR). Similarly, the sCMOS data for Imaging FCS, performed with 100X magnification, was also binned 2×2. The diffusion law plot for sCMOS was performed for square binning of 3 to 7 in contrast to the EMCCD data shown in Fig. 3 where square binning of 1-5 was used. For the N&B analysis using the sCMOS camera, the pixels were 2×2 spatially binned and also time binned to 20 ms. Unless otherwise stated, spatial binning was not performed for SRRF microscopy, but frames were time binned up to 200 ms. All the analyses were performed using the GPU based plugin in ImageJ, described in section 2.7.

#### 1.6 N&B analysis

The time-varying intensity in a pixel (*I*) is determined by the brightness (*B*) of particles and the time-varying number of particles (*N*) in that pixel.

$$\langle I\rangle = \langle N\rangle B \qquad Eq.\ 2$$

where $\langle I\rangle$ is the time-averaged intensity and $\langle N\rangle$ is the time-averaged number of particles. The variance of the intensity is determined only by the variance of the number of particles within the observation volume since the brightness of fluorescent particles is assumed to be a constant within the measurement time. For any random variable *X* and a constant *k*,

$$Var(kX) = k^2 Var(X) \qquad Eq.\ 3$$

Hence

$$\sigma^2 = B^2 Var(N) \qquad Eq.\ 4$$

where $\sigma^2$ is the variance of the fluorescence intensity in the pixel. The number of particles within the observation volume follows a Poissonian distribution. Hence the variance of the time-varying number of particles is equal to the mean of the number of particles within the observation volume.

$$\sigma^2 = B^2\langle N\rangle \qquad Eq.\ 5$$

$$\sigma^2 = B\langle I\rangle \qquad Eq.\ 6$$

Hence the N&B equations are defined as[8]:

$$B = \frac{\sigma^2}{\langle I\rangle} \qquad Eq.\ 7$$

$$N = \frac{\langle I\rangle^2}{\sigma^2} \qquad Eq.\ 8$$

##### 1.6.1 Brightness calibration

The oligomerization state of a protein is determined by computing the ratio of the brightness of the proteins to the monomeric state of the fluorescent protein (FP) used. Not all molecules of fluorescent proteins are fluorescent due to incomplete maturation, misfolding, photo-bleaching issues and possible dark states of the fluorescent moiety[10-12]. Hence in order to estimate the oligomerization state of a protein, one needs to estimate the proportion of FPs that are fluorescent ($p$). The value $p$ is estimated by computing the brightness of two different constructs, monomeric FP and two FPs in tandem (referred to as dimeric FP). Since the proportion of fluorescent FPs is $p$, the proportion of non-fluorescent FPs is *1-p*. Since the dimer sample consists of both fluorescent and non-fluorescent molecules, there are 4 possibilities of brightness: $p^2, p(1-p), (1-p)p$ and $(1-p)^2$.

In a population consisting of $N_1$ monomers, the number of fluorescent molecules are $pN_1$. If the mean brightness of an individual fluorophore is $\varepsilon$, the average intensity $\langle I \rangle_m$ of the population is $\varepsilon p N_1$.

$$\langle I \rangle_m = \varepsilon p N_1 \qquad \textit{Eq. 9}$$

The variance in the number of fluorescent molecules is same as the mean number of fluorescent molecules since the number of fluorescent molecules in the observation volume is Poisson distributed. The variance scales as the square of the brightness. Hence, the variance of the intensity ($\sigma_m^2$) is

$$\sigma_m{}^2 = \varepsilon^2 p N_1 \qquad \textit{Eq. 10}$$

For the dimer sample with the brightness of monomer and dimer being $\varepsilon$ and $2\varepsilon$, respectively, only molecules containing at least one fluorescent FP will contribute to the intensity. Hence the average intensity of the dimer ($\langle I \rangle_d$) of $N_2$ molecules is written as

$$\langle I \rangle_d = 2\varepsilon p^2 N_2 + 2\varepsilon p(1-p)N_2 = 2\varepsilon p N_2 \qquad \textit{Eq. 11}$$

The variance of the intensity of the dimer ($\sigma_d^2$) is given below.

$$\sigma_d{}^2 = (2\varepsilon)^2 p^2 N_2 + 2\varepsilon^2 p(1-p)N_2 = 2\varepsilon^2 p^2 N_2 + 2\varepsilon^2 p N_2 \qquad \textit{Eq. 12}$$

Substituting *Eq. 10*, *Eq. 11* and *Eq. 12*, in *Eq. 7*

$$B_m = \frac{\sigma_m{}^2}{\langle I \rangle_m} = \varepsilon \qquad \textit{Eq. 13}$$

$$B_d = \frac{\sigma_d{}^2}{\langle I \rangle_d} = \varepsilon(p+1) \qquad \textit{Eq. 14}$$

where $B_m$ is the brightness of monomer and $B_d$ is the brightness of the dimer. Dividing *Eq. 14* by *Eq. 13* and rearranging the terms, we obtain an equation to estimate the proportion of fluorescent molecules of an FP:

$$p = \frac{B_d}{B_m} - 1 \qquad \textit{Eq. 15}$$

Denoting the ratio of the brightness of the dimer to the brightness of monomer as $x$, one obtains

$$p = x - 1 \qquad \textit{Eq. 16}$$

##### 1.6.2 Experimentally measured parameters and the error calculations

In the case of N&B analysis, the average brightness of each cell is estimated as an average of brightness of all the pixels within the cell. The data from many such cells were pooled to obtain the population mean and error of brightness of cells.

###### *1.6.2.1 Pooled mean and SEM of brightness*

For each cell, the arithmetic mean and standard deviation (SD) were calculated from the brightness values of pixels in an image of the cell after filtering. Filtering was performed on the *B* map, to ensure that background pixels outside the cells were excluded. A typical threshold of atleast 1500 counts was used. The SD was converted to standard error of the mean (SEM).

$$SD_j = \sqrt{\frac{\sum_{i=1}^{n_j}(B_{i,j} - B_{mean_j})^2}{n_j}} \qquad \textit{Eq. 17}$$

$$SEM_j = \frac{SD_j}{\sqrt{n_j}} \qquad \textit{Eq. 18}$$

where $B_{i,j}$ is the brightness of an individual pixel $i$ in cell $j$, $B_{mean_j}$ is the mean brightness of the pixels in cell $j$, and $n_j$ is the number of pixels in cell $j$. The pooled weighted arithmetic mean brightness and pooled SEM were calculated.

$$B_{pooled} = \frac{\sum_{j=1}^{N} n_j B_{mean_j}}{\sum_{j=1}^{N} n_j} \qquad \textit{Eq. 19}$$

$$SEM_{pooled} = \sqrt{\frac{\sum_{j=1}^{N} SEM_j^2 \; + \; \sum_{j=1}^{N} \frac{\left(B_{mean_j} - B_{pooled}\right)^2}{n_j}}{N}} \qquad \textit{Eq. 20}$$

where $N$ is the total number of cells.

###### *1.6.2.2 Error associated with derived parameters*

In the case of derived parameters, partial derivatives were used to perform propagation on the errors of experimentally measured parameters. For the parameters defined in Sec. 1.6.1,

$$x = \frac{B_{d,pooled}}{B_{m,pooled}} \qquad \textit{Eq. 21}$$

$$\Delta x = x\sqrt{\left(\frac{\Delta B_m}{B_{m,pooled}}\right)^2 + \left(\frac{\Delta B_d}{B_{d,pooled}}\right)^2} \qquad \textit{Eq. 22}$$

where, $\Delta x$ is the propagated SEM of the pooled dimer-to-pooled monomer brightness ratio, $B_d$ is the pooled mean brightness of dimer measurements, $\Delta B_d$ is the pooled SEM of dimer measurements, $B_m$ is the pooled mean brightness of monomer measurements, and $\Delta B_m$ is the pooled SEM of monomer measurements.

The proportion of fluorescent molecules of an FP is obtained by subtracting a value of 1 from the ratio of the brightness of the dimer to the brightness of the monomer. Hence the error associated with the proportion of FPs being fluorescent is

$$\Delta p = \Delta x \qquad \textit{Eq. 23}$$

where $\Delta p$ is the SEM of the proportion of FPs which are fluorescent.

##### 1.6.3 Brightness of EGFR

The EGFR sample has a mixture of both monomers and dimers. The overall intensity is the sum of the contributions from the monomer and the dimer (see *Eqs. 9* and *11*).

$$\langle I \rangle_E \; = \; \varepsilon p N_1 + 2\varepsilon p N_2 \qquad \textit{Eq. 24}$$

The overall variance of intensity of EGFR is the sum of the variances of the intensities of the monomer and dimer components (*Eqs. 10* and *12*). Hence,

$$\sigma_E{}^2 = \varepsilon^2 p N_1 + 2\varepsilon^2 p^2 N_2 + 2\varepsilon^2 p N_2 \qquad \textit{Eq. 25}$$

where $\langle I \rangle_E$is the average intensity of EGFR sample and $\sigma_E{}^2$ is the variance of the intensity of EGFR sample. Substituting *Eq. 25* and in *Eq. 7*, and simplifying the equation,

$$B_E = \frac{\varepsilon(N_1 + 2pN_2 + 2N_2)}{N_1 + 2N_2} \qquad \textit{Eq. 26}$$

where $B_E$ is the brightness of EGFR. Dividing *Eq. 26* by *Eq. 13*

$$\frac{B_E}{B_m} = r = \frac{N_1 + 2pN_2 + 2N_2}{N_1 + 2N_2} \qquad \textit{Eq. 27}$$

where $r$ is the ratio of the brightness of EGFR to the brightness of monomer. If the monomer EGFR population fraction is $f$ and dimer EGFR population fraction is $1\text{-}f$,

$$f = \frac{N_1}{N_1 + N_2} \qquad \textit{Eq. 28}$$

$$f = \frac{2(p - r + 1)}{2p - r + 1} \qquad \textit{Eq. 29}$$

$$1 - f = e_d = \frac{r - 1}{2p - r + 1} \qquad \textit{Eq. 30}$$

where $e_d$ is the dimer fraction. Rewriting the EGFR dimer fraction in terms of EGFR monomers present as dimers,

$$m_e = \frac{2e_d}{e_d + 1} \qquad \textit{Eq. 31}$$

where $m_e$ is the mean fraction of EGFR monomers present as dimers.

###### *1.6.3.1 Error calculations of derived parameters*

Using partial derivatives, the error associated with the derived parameters in Sec. 1.6.3 were determined.

$$\Delta r = \frac{B_E}{B_m}\sqrt{\left(\frac{\Delta B_m}{B_m}\right)^2 + \left(\frac{\Delta B_E}{B_E}\right)^2} \qquad \textit{Eq. 32}$$

where $\Delta r$ is the propagated SEM of the EGFR-to-monomer brightness ratio, $B_E$ is the pooled mean brightness of EGFR measurements, and $\Delta B_E$ is the pooled SEM of EGFR measurements.

$$\Delta e_d = \frac{2}{(2p - r + 1)^2}\sqrt{p^2\Delta r^2 + (r-1)^2\Delta p^2} \qquad \textit{Eq. 33}$$

where $\Delta e_d$ is the SEM of the EGFR dimer fraction.

$$\Delta m_e = \frac{2\Delta e_d}{(e_d + 1)^2} \qquad \textit{Eq. 34}$$

where $\Delta m_e$ is the SEM of this fraction.

##### 1.6.4 Brightness of oligomers

In the case where the sample contains oligomers of order '*n*'

$$\langle I \rangle_n = N\sum_{i=1}^{n}(i\varepsilon)\binom{n}{i}p^i(1-p)^{n-i} \qquad \textit{Eq. 35}$$

$$\sigma_n{}^2 = N\sum_{i=1}^{n}(i\varepsilon)^2\binom{n}{i}p^i(1-p)^{n-i} \qquad \textit{Eq. 36}$$

where $\sigma_n{}^2$ is the variance of the intensity of the oligomer, $\langle I \rangle_n$ is the average intensity of oligomer sample, $N$ is the number of oligomer molecules. Substituting, the variance and intensity in *Eq. 7*

$$B_n = \frac{\sum_{i=1}^{n}(i\varepsilon)^2\binom{n}{i}p^i(1-p)^{n-i}}{\sum_{i=1}^{n} i\varepsilon\binom{n}{i}p^i(1-p)^{n-i}} \qquad \textit{Eq. 37}$$

$$B_n = \varepsilon[(n-1)p + 1] \qquad \textit{Eq. 38}$$

where $B_n$ is the brightness[11] of oligomer of order *n*. Dividing by *Eq. 13*,

$$\frac{B_n}{B_m} = (n-1)p + 1 \qquad Eq.\ 39$$

##### *1.6.4.1 Estimation of EGFR oligomerization*

For an EGFR sample containing a mixture of oligomers up to an oligomer of order $n$,

$$B_{E,oligo} = \frac{\sum_{i=1}^{n} N_i \sum_{j=1}^{i} (j\varepsilon)^2 \binom{i}{j} p^j (1-p)^{i-j}}{\sum_{i=1}^{n} N_i \sum_{j=1}^{i} j\varepsilon \binom{i}{j} p^j (1-p)^{i-j}} \qquad Eq.\ 40$$

where $B_{E,oligo}$ is the brightness of the EGFR in oligomeric state, $N_i$ is the number of oligomer molecules of order-$i$.

Assuming a mix of EGFR monomers, dimers and trimers only, where $N_1$, $N_2$ and $N_3$ are the number of monomers, dimers and trimers respectively,

$$B_{E,trimer} = \frac{\varepsilon(N_1 + 2N_2 + 2pN_2 + 3N_3 + 6pN_3)}{N_1 + 2N_2 + 3N_3} \qquad Eq.\ 41$$

where $B_{E,trimer}$ is the brightness of the EGFR in trimeric state.

The ratio between brightness of EGFR trimer to that of the monomer is

$$r_{trimer} = \frac{B_{E,trimer}}{B_{monomer}} = \frac{N_1 + 2N_2 + 2pN_2 + 3N_3 + 6pN_3}{N_1 + 2N_2 + 3N_3} \qquad Eq.\ 42$$

where $r_{trimer}$ is the ratio of the brightness of the EGFR trimer to the brightness of the monomer.

Let $m$, $d$ and $t$ be the fraction of monomers, dimers and trimers, respectively.

$$m = 1 - d - t = \frac{N_1}{N_1 + N_2 + N_3} \qquad Eq.\ 43$$

$$d = \frac{N_2}{N_1 + N_2 + N_3} \qquad Eq.\ 44$$

$$t = \frac{N_3}{N_1 + N_2 + N_3} \qquad Eq.\ 45$$

$$r_{trimer} = \frac{1 + d + 2pd + 2t + 6pt}{1 + d + 2t} \qquad Eq.\ 46$$

Rewriting $[m;\ d;\ t]$ in terms of $r_{trimer}$

$$\left[1 - d - t; d; \frac{1 + d + 2pd - r_{trimer}(1+d)}{2r_{trimer} - 2 - 6p}\right]$$

If no dimers:

$$\left[1-t;0;\frac{r_{trimer}-1}{6p-2r_{trimer}+2}\right]$$

If no trimers:

$$\left[1-d;\frac{r_{trimer}-1}{2p-r_{trimer}+1};\ 0\right]$$

It is important to note that a complete solution can be obtained only for two oligomer species. Hence the solutions shown above indicate special cases where either the trimer or the dimer is absent. If there are more than two kinds of oligomers, complete solutions are possible only with when more measurement parameters are available. Therefore, we provide only apparent oligomer percentages with the understanding that they represent a minimal model of monomers and one oligomer species that are consistent with the experimental data. We can however, not exclude that the sample consists of a mixture of oligomers including higher oligomer species.

#### 1.7 GPU based plugin

The input for the Imaging FCS plugin in ImageJ is a 16-bit tiff image stack. Autocorrelations are calculated at every pixel, and fitted with an appropriate model. The GPU code parallelizes the calculation of the correlation functions and their fits.

The described GPU acceleration has been performed for the computation of autocorrelations, N&B analysis and computation of FCS diffusion laws. In the case of the computation of autocorrelations, the binning, bleach correction, and calculation of the correlations are all performed on the GPU. The fitting is performed in the GPU using the open source GPUfit program[13]. The flow chart showing the sequence of calculations is shown in Fig. S1a.

In order to run calculations on an NVIDIA GPU installed in a machine with a 64-bit Windows or Linux operating system, *agpufit.dll* or *libagpufit.so* is loaded in Windows or Linux respectively upon running Imaging FCS to check for availability of CUDA runtime environment using *isCudaAvailable()*.

If the criteria are not met, the program will perform calculations on the CPU instead. The input to the Imaging FCS plugin is a stack of images of dimensions [*r*, *c*, *t*] where *r*, *c* and *t* are the number of rows, columns and timepoints respectively. The bridge from Imaging FCS to CUDA code, which are written in C++, is made possible through the Java Native Interface (JNI). The JNI static functions are coded in *gpufitImFCS.java*.

##### 1.7.1 GPU Kernel

Depending upon the user parameters, the entire or a part of the image stack of dimensions [*win*, *hin*, t] is transferred to the GPU. ACF calculation on the GPU is parallelized along each pixel of the image. The parameters *wtemp* and *htemp* (Ref. Fig. S1 for definitions) determines CUDA kernel size, i.e. the number of blocks per grid, which is denoted with prefix *gridSize* in the CUDA code. The number of threads per block, which is denoted with prefix *blockSize*, is set at 16 x 16 x 1. It was found that maximising *blockSize* at 32 x 32 x 1 may sometimes result in the error ‘too many resources requested’, which is the reason for our choice of a smaller *blockSize*. On the same token, parameter *power_of_two_n_points* in *Info::configure()* function is increased by a factor of 4 in comparison to the original *gpufit* code because of the complexity of the *ACF_1D CUDA* kernel fitting function.

##### 1.7.2 Array sizes

The JNI API *SetFloatArrayRegion* has limited output array sizes. We therefore limited 2×2 binning of stacks 50,000 frames to 96 pixels × 96 pixels. Binning of larger stacks, especially full frames of 128 pixels × 128 pixels × 50,000 frames, are binned by the CPU. This is not an intrinsic limit of the method, but a technical issue of the GPU memory size. If an error is encountered while doing a calculation in GPU mode, the program will then perform the calculation on CPU.

##### 1.7.3 Comparison of processing time between CPU and GPU

Random walk simulations as described here[2] were used to simulate a dataset which was used for estimating the processing time using a CPU and GPU.

##### 1.7.4 Program availability

The plugin is available for download using an upload site in ImageJ and is also available at http://www.dbs.nus.edu.sg/lab/BFL/imfcs_image_j_plugin.html or can be obtained within ImageJ from the ImageJ update sites.

#### 2 Supplementary figures and tables

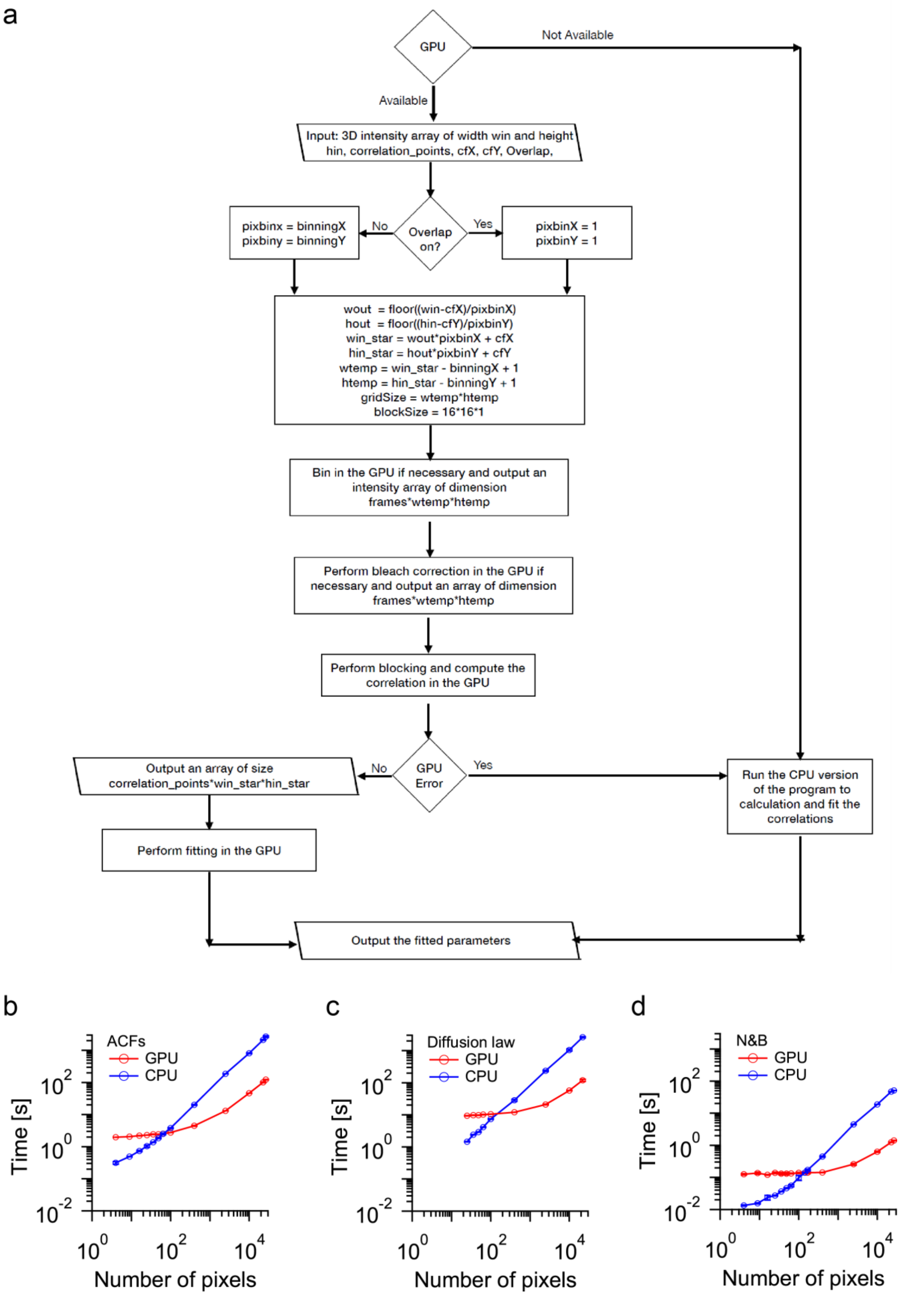


**Fig. S1: GPU based processing**: **(a)** Flow chart describing the flow of control on the GPU, and the computation time comparison between GPU and CPU for **(b)** ACFs fitting, **(c)** diffusion law and **(d)** N&B

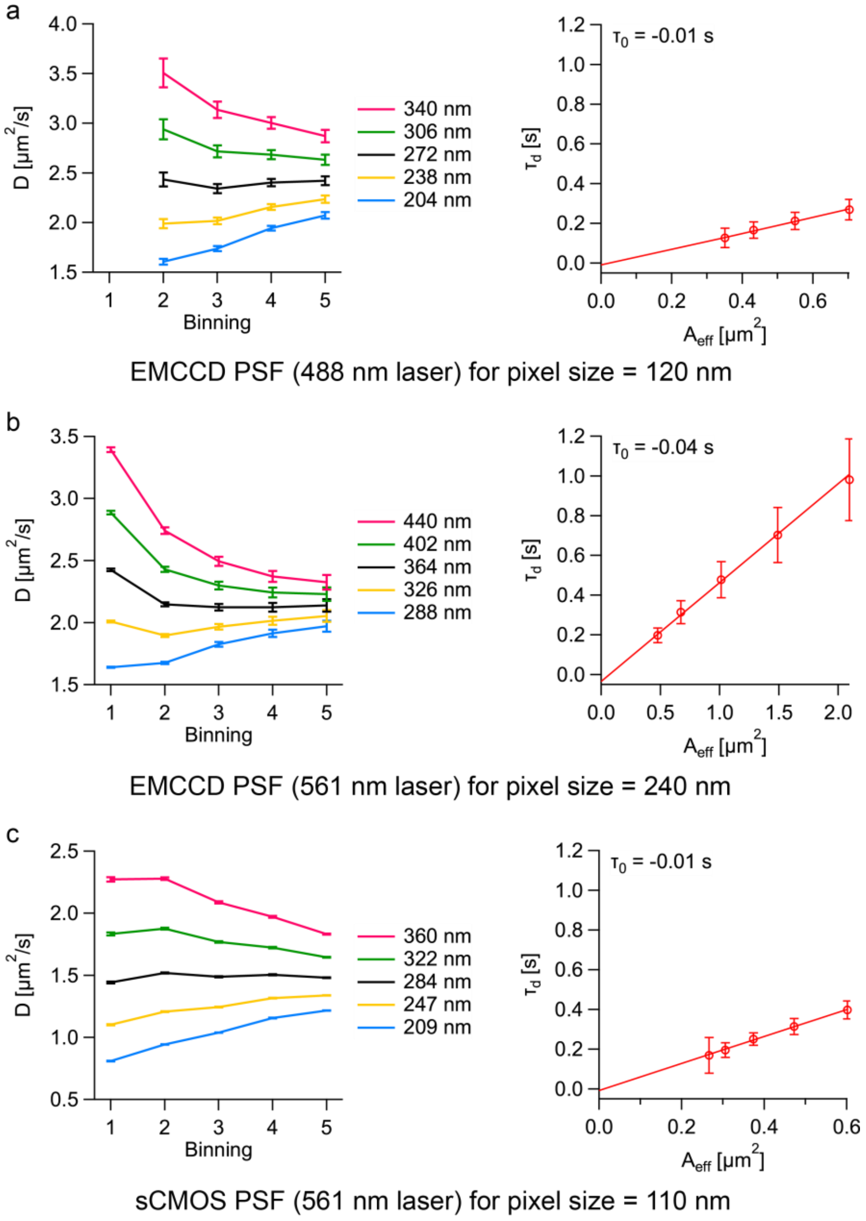


**Fig. S2: PSF calibration and diffusion law plots**: The PSF calibration and diffusion law plots for **(a)** the EMCCD camera at a magnification of 200X (120 nm pixel size in object space) at a wavelength of 488 nm, **(b)** the EMCCD camera at a magnification of 100X (240 nm pixel size in object space) at a wavelength of 561 nm, and **(c)** the sCMOS camera at a magnification of 100X (110 nm pixel size in object space) at a wavelength of 561 nm are shown here. The detailed experimental configurations are provided in Table S2.

**Table S2: *1/e²* radii of various experimental configurations**

| | Laser [nm] | Camera | $P_{chip}$ [μm] | Obj | Additional magnification | $P_{mag}$ [nm] | PSF [nm] | $D_{ave}^{*}$ [μm²/s] |
|---|---|---|---|---|---|---|---|---|
| **(a)** | **488** | EMCCD | 24 | 100X | 2X | 120 | 272 | 1.97 ± 0.91 |
| **(b)** | **561** | EMCCD | 24 | 100X | None | 240 | 364 | 2.00 ± 0.34 |
| **(c)** | **561** | sCMOS | 11 | 100X | None | 110 | 284 | 1.79 ± 0.39 |

**$P_{chip}$**-Pixel size on chip; **$P_{mag}$**-Pixel size after magnification; **Obj**-Objective;

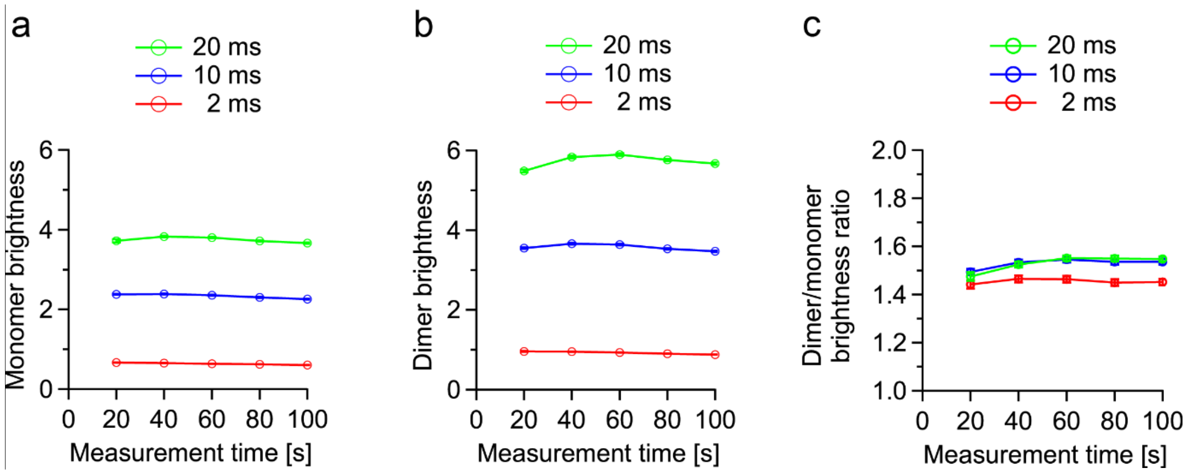


**Fig. S3: Number and Brightness analysis – Effect of exposure time and measurement time on brightness. (a)** The mean brightness of PMT-mApple is plotted against the total measurement time, for different exposure times (2 ms, 10 ms, 20 ms). **(b)** The mean brightness of PMT-mApple$_2$ is plotted against the total measurement time, for different exposure times (2 ms, 10 ms, 20 ms). **(c)** The brightness ratio of PMT-mApple$_2$/PMT-mApple is plotted against total measurement time, for different exposure times (2 ms, 10 ms, 20 ms). Each point is an average of 3 different cell measurements for both PMT-mApple and PMT-mApple$_2$. The number of pixels in each cell at each acquisition time is different due to differential intensity filtering in each case, but each cell had at least 1850 and at most 6250 valid pixels.

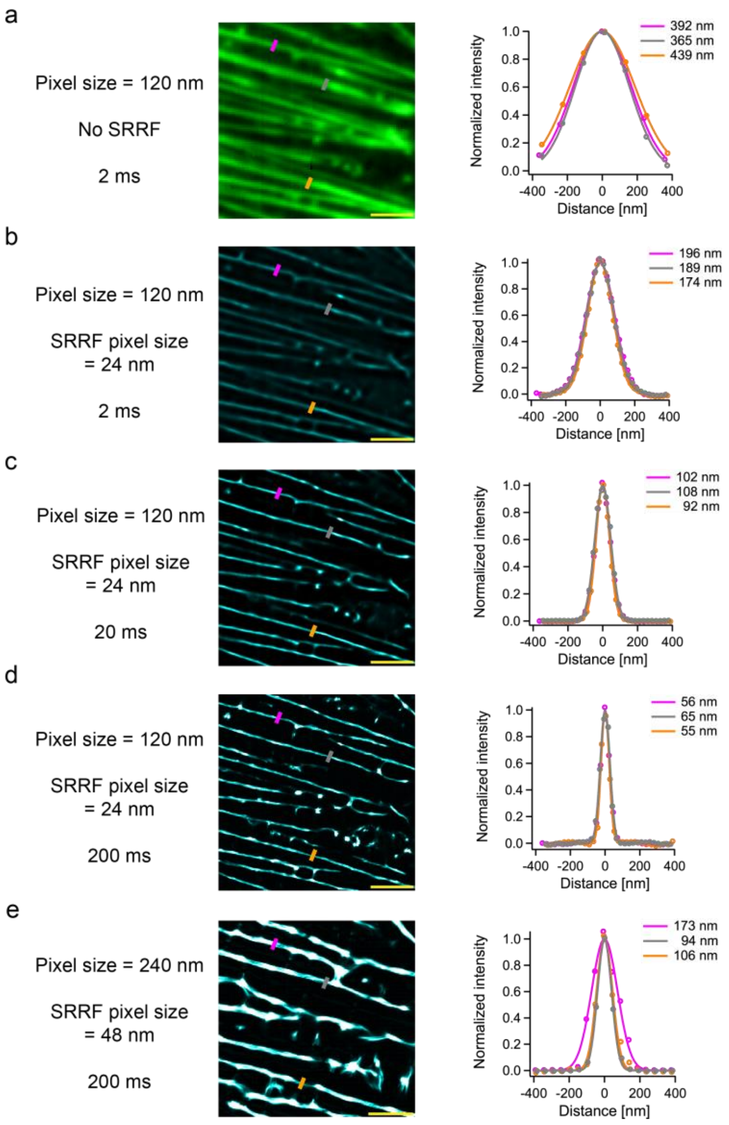


**Fig. S4: Time binning and magnification improves spatial resolution in SRRF:** In panels (a)-(e), images from TIRF or SRRF as indicated are shown on the left. Normalized intensity profiles across actin fibres at three different positions (indicated by magenta, grey, and orange lines on the cell image) are shown on the right. All the reported values in the figure are the FWHM of Gaussian fits to the intensity profiles **(a**) TIRF image of a CHO-K1 cell labeled with LifeAct-EGFP at 200X magnification. **(b)** SRRF map of the 2 ms data shown in image (a). **(c)** and **(d)** show the SRRF maps after time binning of the 2 ms data shown in image (a) to 20 ms and 200 ms, respectively. **(e)** SRRF map and histogram generated after time binning of the 2 ms data of the same area at 100X magnification to 200 ms. The FWHM (mean ± SD) of Gaussian Fits to the normalized intensity profiles from each image is as follows: (a) 399 ± 31 nm, (b) 186 ± 9 nm, (c) 100 ± 6 nm, (d) 59 ± 5 nm, and (e) 124 ± 35 nm. The pixel sizes reported are after magnification (refer Table 2). The scale bars shown in yellow measure 2.5 µm in images (a)-(e).

#### Table S3: FRC and P2P measures of resolution

| Camera | Pixel size in SRRF [nm] | Mean FRC [nm] | P2P [nm] | Area on SRRF map | Based on figure |
|---|---|---|---|---|---|
| **EMCCD** | 48 nm | 145.3 ± 18.3 | 192 | 200 x 200 | 1e* |
| **EMCCD** | 24 nm | 77.6 ± 21.1 | 120 | 460 x 460 | S5d |

*P2P map not shown

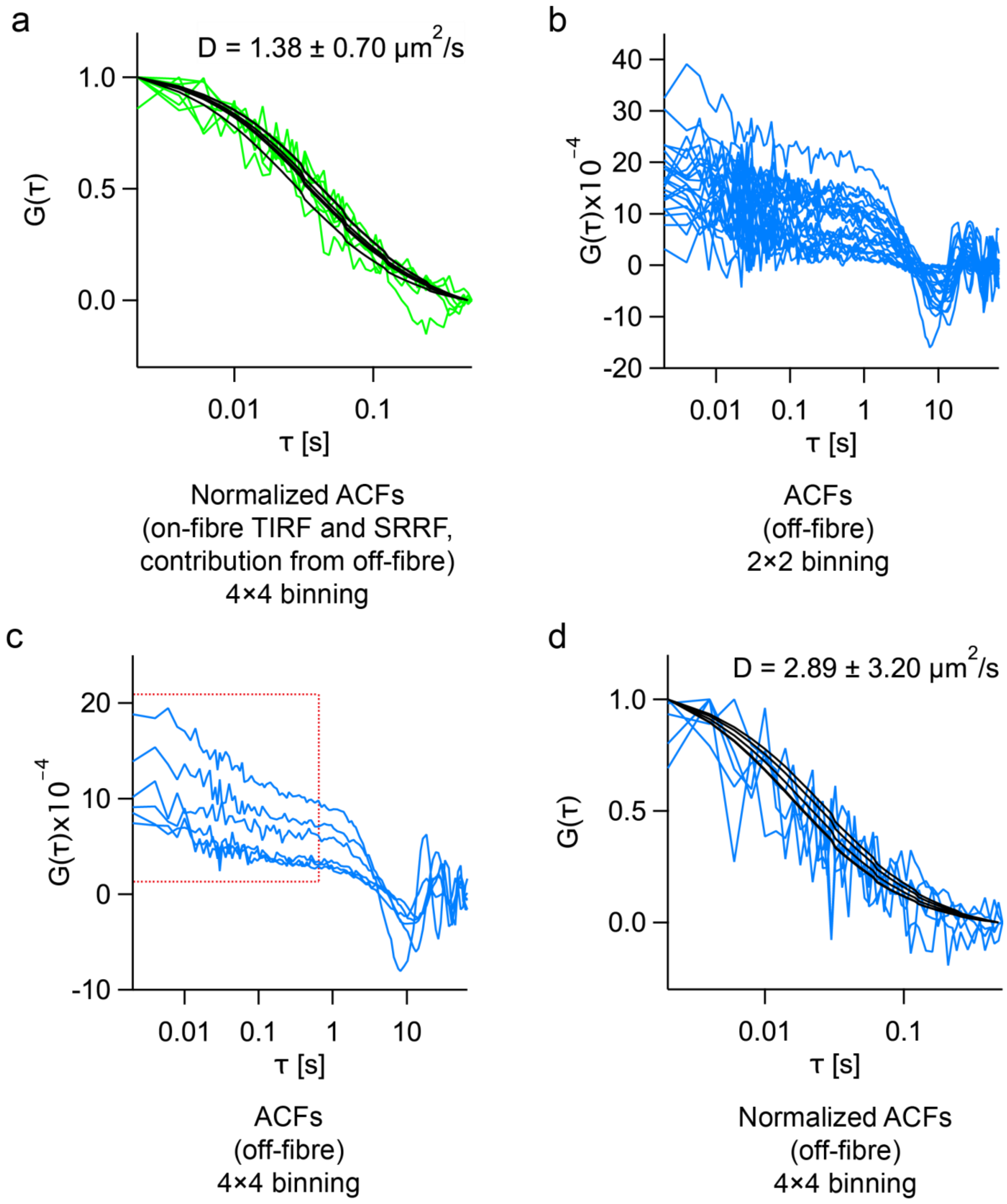


**Fig. S5: Off-fibre areas show a fast diffusion component: (a)** Normalized ACFs of LifeAct-EGFP from fibre areas (on-fibre TIRF and SRRF with contribution from off-fibre; Fig. 2f-case 5) after 4×4 binning at 200X magnification. **(b)** ACFs of LifeAct-EGFP from off-fibre areas (Fig. 2f-case 4) after 2×2 binning at 200X magnification **(c)** ACFs of LifeAct-EGFP from the same areas in (b) after 4×4 binning (Fig. 2f-case 6) at 200X magnification. The fast component is marked with a red box. **(d)** Normalized ACFs of the fast diffusion component. Mean ± SD.

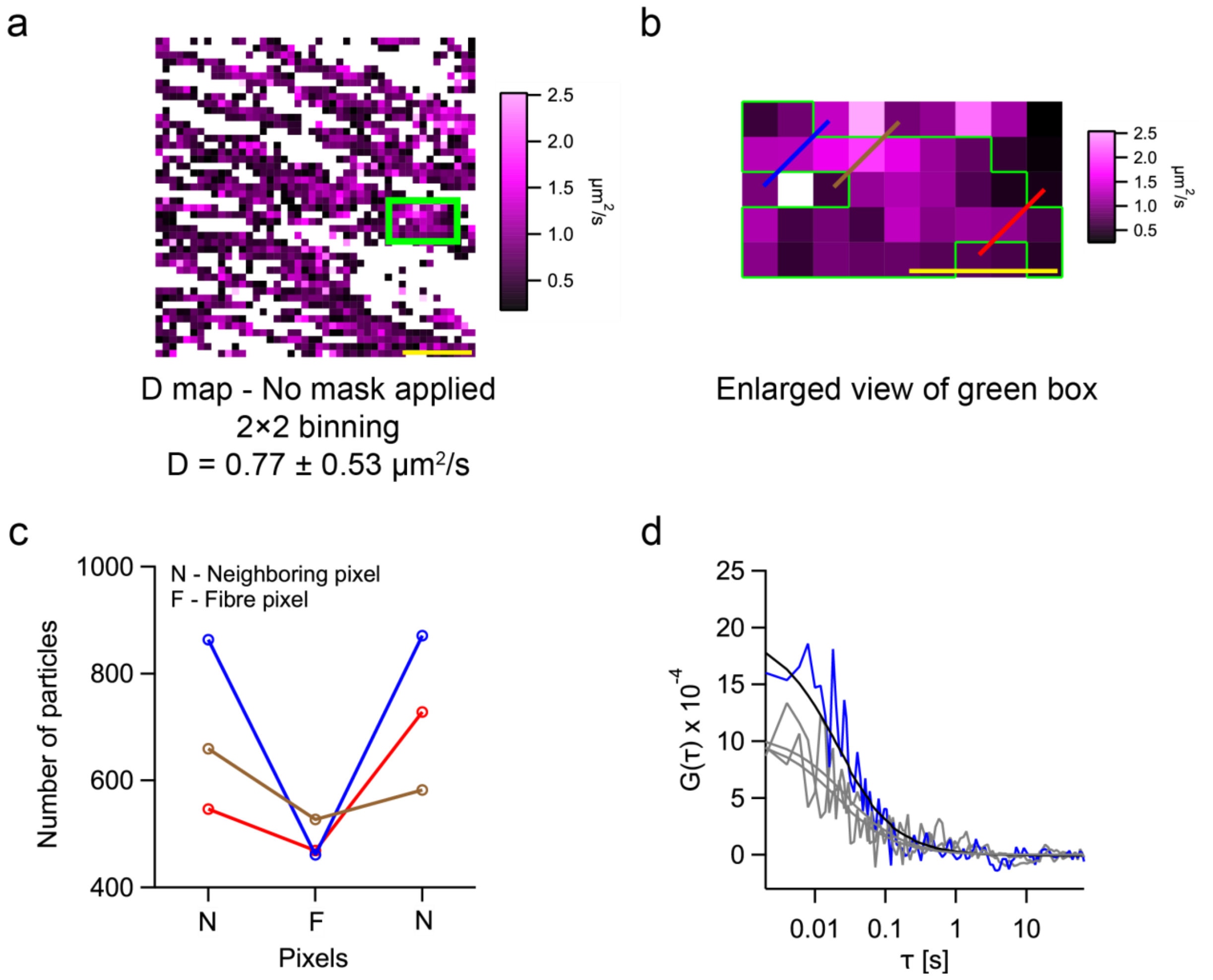


**Fig. S6: SRRF filtered *D* map: (a)** The *D* map generated after thresholding (0.2-100 µm²/s; *D* = 0.77 ± 0.53 µm²/s)(Fig. 2f-case 2). The coefficient of variation (COV-ratio of standard deviation to the mean) is 69%. **(b)** The enlarged view of the area labeled with the green box in (a). The pixels bounded by the green lines are the pixels that overlap with SRRF map (white pixels in Fig. 2d). **(c)** Plot of number of particles versus the pixel type (F: fibre, N: non-fibre) for the three colored lines drawn in (b). Each colored line covers 3 pixels – a central pixel (on-fibre SRRF and TIRF, Fig. 2f-case1) flanked by two neighboring pixels (on-fibre TIRF, off-fibre SRRF, contribution from off-fibre, Fig. 2f-case3). All the pixels are plotted from the left to right for each line. **(d)** ACFs of the blue line in (b) shown as an example. The ACF of the on-fibre SRRF and TIRF pixel is shown in blue, while the ACFs of the neighboring pixels are shown in grey.

#### Table S4: *N* and *D* values for fibre and non-fibre areas

| Image | *N* per pixel | *D* [µm²/s] | Pixels |
|---|---|---|---|
| **Whole image** | 912 ± 523 | 0.77 ± 0.53 | 1247 |
| **Fibres only** | 685 ± 221 | 0.80 ± 0.39 | 670 |
| **Off-fibre only** | 1176 ± 637 | 0.77 ± 0.66 | 577 |

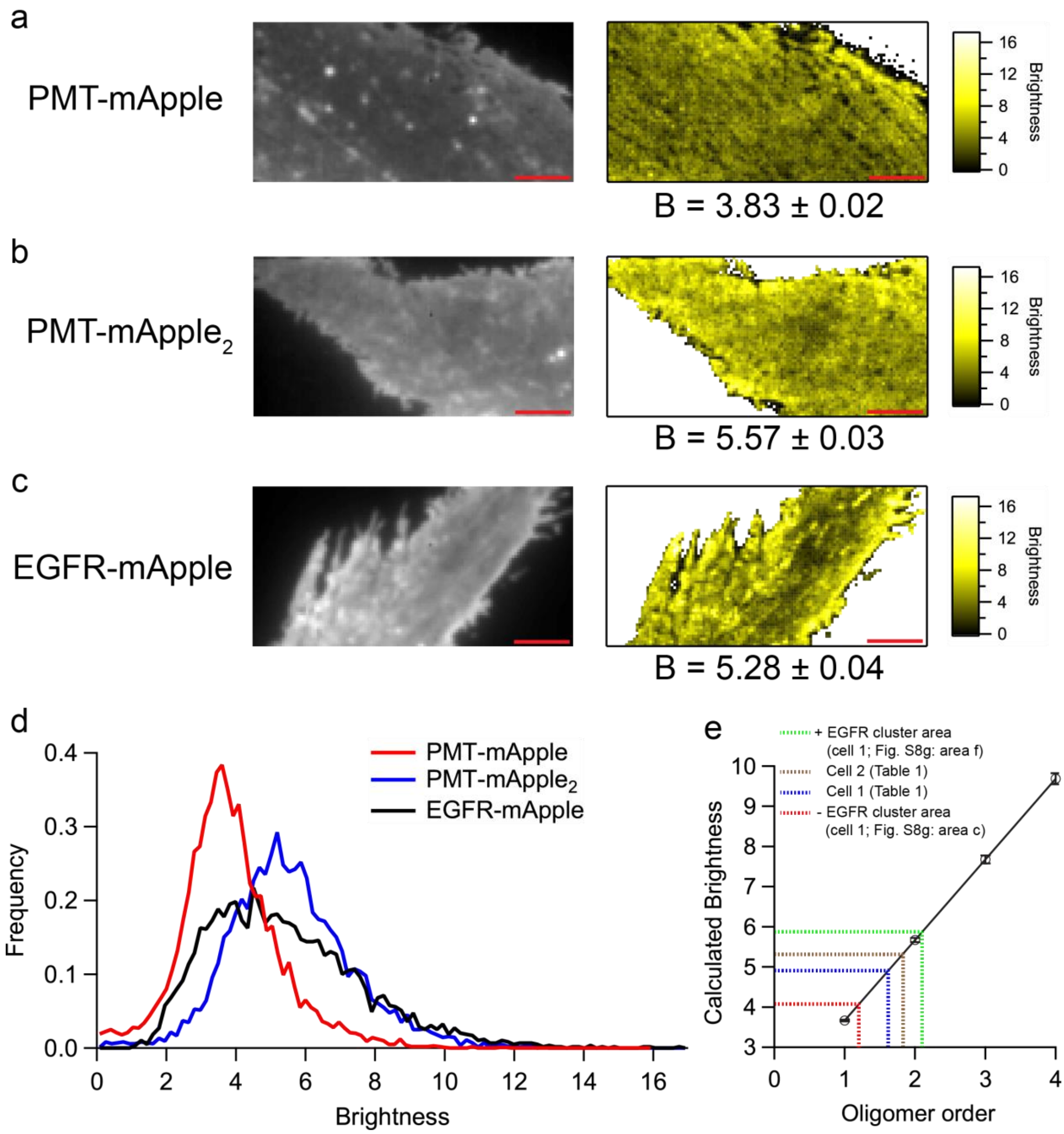


**Fig. S7: Number and Brightness analysis. (a)** TIRF (left) and brightness (right) images of CHO-K1 cells expressing PMT-mApple, PMT-mApple$_2$ and EGFR-mApple (cell 2 in Table 1) are shown in **(a)**, **(b)** and **(c)**, respectively. The brightness histograms corresponding to the images shown in (a)-(c) are shown in **(d)**. **(e)** From the experimentally estimated monomer brightness and probability, the scaling of expected brightness with oligomer order in shown. The dotted lines indicate the average B values for different cases – an area with EGFR cluster in cell 1 (green; refer Fig. S8g: area f), cell 2 (brown; refer Table 1), cell 1 (blue; refer Table 1) and an area without visible EGFR clusters in cell 1 (red; refer Fig. S8g: area c). The scale bars shown in red measure 5 µm in images (a)-(c). The reported values in the figure are mean ± SEM.

**Table S5: Diffusion law intercepts in different areas of CHO-K1 cells labeled with EGFR-mApple***

| Cell | Diffusion law intercept [s] | | | |
|---|---|---|---|---|
| | + Actin fibre<br><br>+ EGFR cluster | + Actin fibre<br><br>- EGFR cluster | - Actin<br><br>+ EGFR cluster | - Actin fibre<br><br>- EGFR cluster |
| 1 | 2.56 | 2.01 | 1.81 | 1.81 |
| 2 | 2.53 | 2.42 | 3.43 | - |
| 3 | 2.21 | 1.48 | - | 0.40 |
| 4 | 5.3 | 2.4 | 4.47 | 3.16 |
| 5 | 2.02 | 1.37 | - | 1.22 |
| 6 | 2.17 | 2.06 | 1.73 | - |
| **Mean** | **2.80 ± 1.24** | **1.96 ± 0.45** | **2.86 ± 1.33** | **1.65 ± 1.16** |

*Please refer to Table 1 for other corresponding parameters of the cells. Some cells do not contain all types of areas, and thus no values are provided.

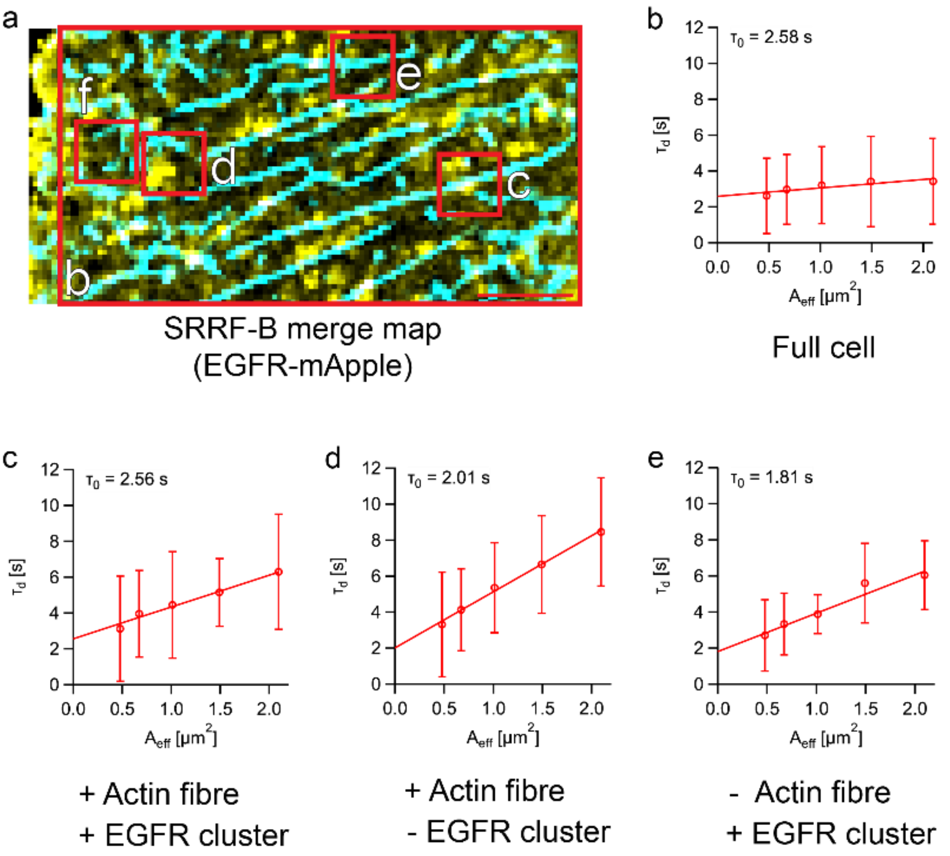


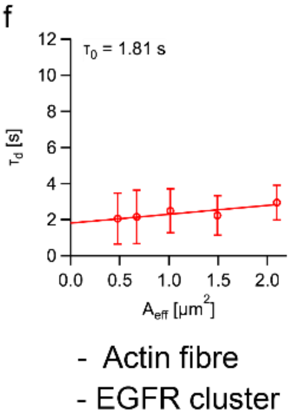


g

| Area | D (μm²/s) | B (dimer fraction) | $T_0$ (s) |
|---|---|---|---|
| b (114×56 pixels) | 0.18 ± 0.15 | 4.80 ± 0.03 (0.57 ± 0.07) | 2.58 |
| c (10×10 pixels) | 0.11 ± 0.08 | 5.87 ± 0.26 (NA) | 2.56 |
| d (10×10 pixels) | 0.16 ± 0.14 | 4.15 ± 0.12 (0.24 ± 0.06) | 2.01 |
| e (10×10 pixels) | 0.18 ± 0.13 | 7.55 ± 0.73 (NA) | 1.81 |
| f (10×10 pixels) | 0.24 ± 0.16 | 4.03 ± 0.11 (0.18 ± 0.06) | 1.81 |

**Fig. S8: Diffusion law analysis over different areas of EGFR-mApple cell: (a)** SRRF-B merged image of a CHO-K1 cell expressing EGFR-mApple (cell 1 from Table 1). The red boxes (numbered b-f) indicate the different areas tested. **(b)** Diffusion law plot in the area marked by the red box b. The intercept is 2.58 s. This is the largest rectangular box (114x56 pixels) that could be fitted into the cell. **(c)** Diffusion law in area marked by the red box c. The intercept is 2.56 s. This area (10x10 pixels) was chosen as there were both an EGFR cluster and an actin fibre. **(d)** Diffusion law in the area marked by the red box d. The intercept is 1.81 s. This area was chosen as there was neither an actin fibre nor an EGFR cluster. **(e)** Diffusion law in the area marked by the red box e. The intercept is 2.01 s. This area (10x10 pixels) was chosen as there was an actin fibre but no EGFR cluster was present. **(f)** Diffusion law in the area marked by the red box f. The intercept is 1.81 s. This area (10x10 pixels) was chosen as there was an EGFR cluster but no actin fibre was present. **(g)** Table of *D* (mean ± SD), *B* (dimer fraction in brackets; mean ± SEM), and intercept values in the chosen areas (b)-(f). For *B* higher than dimer control, the dimer model is not applicable and an oligomer model has to be used. The regions with cluster have a higher brightness when compared to those which do not have clusters as seen in Table g. The scale bar in red measures 5 μm in image (a).

### 3 Supplementary results: Use of sCMOS in multi-parametric analysis

Here, we describe the optimization and results obtained using sCMOS as a detector for dual channel measurements on cells labeled with EGFR-mApple and LifeAct. As the fast acquisition limits the photon counts, large pixels or pixel binning are used to optimize the FCS signal. Here we used 2×2 binning on data sets acquired at 2 ms using sCMOS. In the case of diffusion law analysis, it was calculated from 3×3 to 7×7 binning. N&B was performed with 2×2 spatial binning at 20 ms time binning using 5,000 frames.

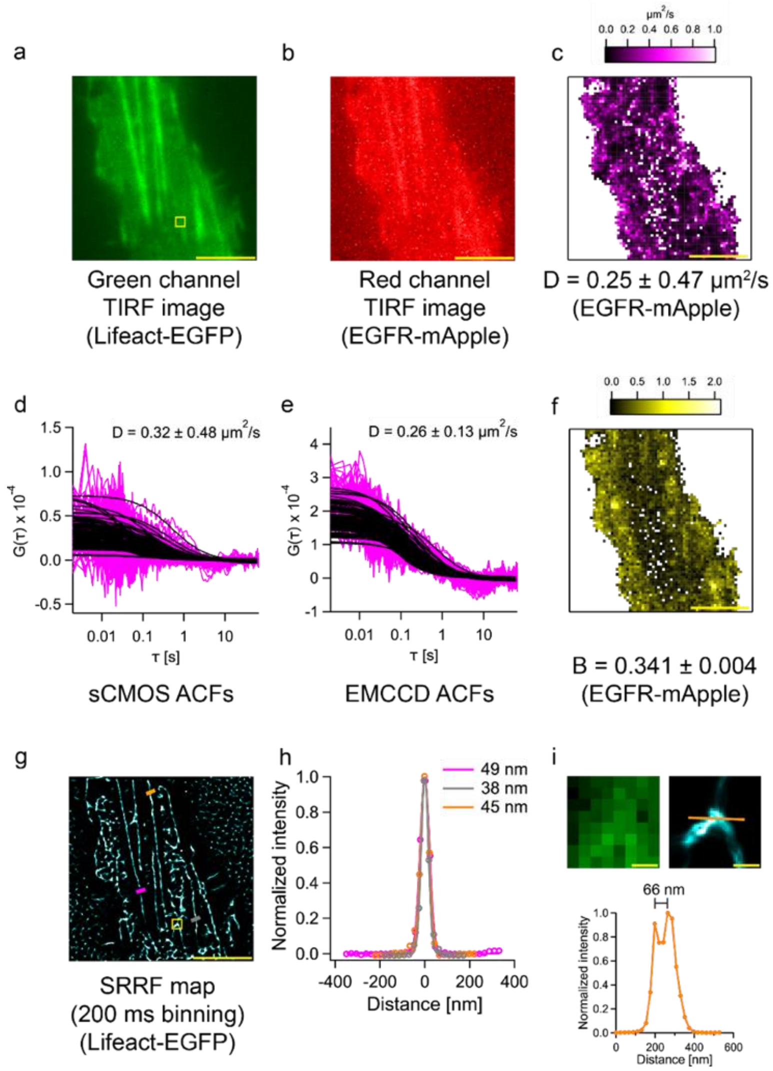


**Fig. S9: Multi-parametric analysis using sCMOS: (a)** and **(b)** are TIRF images of CHO-K1 cells labeled with LifeAct-EGFP and EGFR-mApple. **(c)** Diffusion map of EGFR-mApple **(d)** and **(e)** show ACFs for EGFR-mApple from sCMOS and EMCCD, respectively. The sCMOS (magnified pixel size = 110 nm) data was binned 2×2 (chosen area = 2.2 µm x 2.2 µm), while the EMCCD (magnified pixel size = 240 nm) data was not spatially binned (chosen area = 2.4 µm x 2.4 µm). The mean and standard deviation of the diffusion coefficient estimated only from the displayed curves is shown. **(f)** Brightness map of EGFR-mApple (*B* = 0.341 ± 0.004, dimer fraction = 0.77 ± 0.13; mean ± SEM). **(g)** SRRF map (200 ms binning) of LifeAct EGFP. **(h)** Normalized intensity profile across actin fibres at the three different points on the cell image (e). All the reported values in the figure are the FWHM of the Gaussian fits (average = 44 ± 5 nm). **(i)** Enlarged views of the yellow boxes in images (a) and (g). The intensity profile below shows the actin fibre branching point (indicated by the orange line on the enlarged SRRF image). The peak-to-peak resolution is 66 nm at this point. The scale bars shown in yellow in (a,b,c,f,g) measure 5 µm, and 250 nm in (i). Unless otherwise stated, all values are given as mean ± SD.

The estimated diffusion coefficient of EGFR measured using a sCMOS is 0.25 ± 0.47 μm$^2$/s. This corresponds to a COV of 188%. The COV of diffusion coefficient of EGFR measuring using an EMCCD is 79% (0.19 ± 0.15 μm$^2$/s, Fig. 3, S9e). The COV of sCMOS is larger than that of EMCCD due to a lower SNR of the obtained autocorrelation curves using a sCMOS (Fig. S9). It is important to note that the inclusion of an optosplit in the detection path for two channel measurements leads to additional losses in collected fluorescence signal. The reduction in collected signal intensity leads to lower SNR of ACFs. In the case of measurements with sCMOS, the dimer fraction is found to be 77 ± 13%. The larger error for the dimer fraction of EGFR using the sCMOS camera is attributed to the reduced SNR (Fig. S9). In the case of SRRF, a FWHM of 44 ± 5 nm is obtained using 200 ms binning. The mean FRC was 73.0 ± 16.2 while the P2P is 66 nm (Fig. S9). However, one can clearly detect artefacts outside the cell where no features are expected (Fig. S9e). Hence, we investigated the effects of spatial and temporal binning in reducing the artefacts seen outside the cell perimeter.

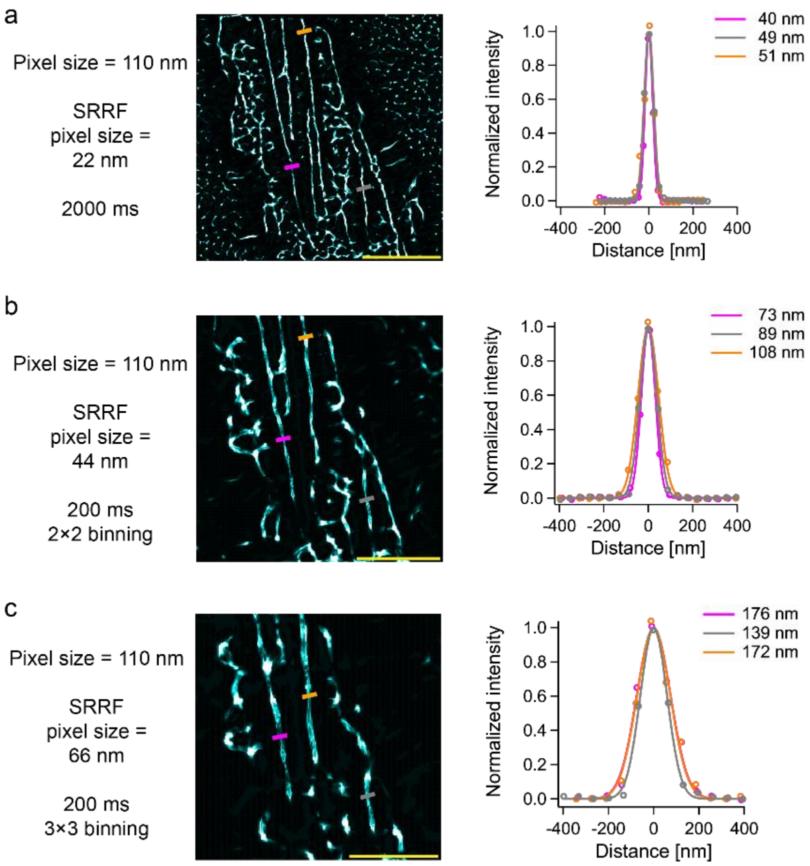


**Fig. S10: Spatial binning of pixels on sCMOS reduces background artefacts in SRRF (a)** SRRF map and normalized intensity profile generated after time binning of the 2 ms data shown in Fig. S9a to 2,000 ms. The normalized intensity profile on the right shows the actin fibre thickness at three different points (indicated by magenta, grey, and orange lines on the cell image on the left). The background artefacts were not reduced. Also, no improvement in the thickness of the actin fibre (refer Fig. S9g) was observed. **(b)** SRRF map and normalized intensity profile generated after 2×2 spatial binning of the pixels in the 2 ms data shown in Fig. S9a. The spatial binning reduced the background artefacts significantly. **(c)** SRRF map and normalized intensity profile generated after 3×3 spatial binning of the pixels in the 2 ms data shown in Fig. S9a. The higher spatial binning further reduced the artefacts. All the reported values in the figure are Gaussian fits to the intensity profiles. The FWHM (mean ± SD) of Gaussian Fits to the normalized intensity profiles for each image is as follows: (a) 47 ± 5 nm, (b) 90 ± 14 nm, and (c) 162 ± 17 nm. The pixel sizes reported are after magnification (refer Table S1). The scale bars shown in yellow measure 5 μm in images (a)-(c).

Increasing the time bin to 2,000 ms did not lead to a removal of artefacts (Fig. S10a). By contrast, increasing the spatial bin to 2×2 and 3×3 led to reduction in artefacts (Fig. S10b and c). But the FWHM of the fibre increased with increased spatial binning. A comparison between Fig. S5 and Fig. S10 illustrates the typical trade-offs in a SRRF investigation. At optimal SNR, a decrease in pixel size leads to an improvement in the FWHM for the fibre (Fig. S4). At sub-optimal SNR, a decrease in pixel size post acquisition through binning leads to an improvement in FWHM but with an increase in the presence of artefactual structural features as well (Fig. S10).

The diffusion law analysis performed in sCMOS is shown in Fig. S11. The intercept values are similar to those obtained using EMCCD as a detector (Table S5, Fig. S8).

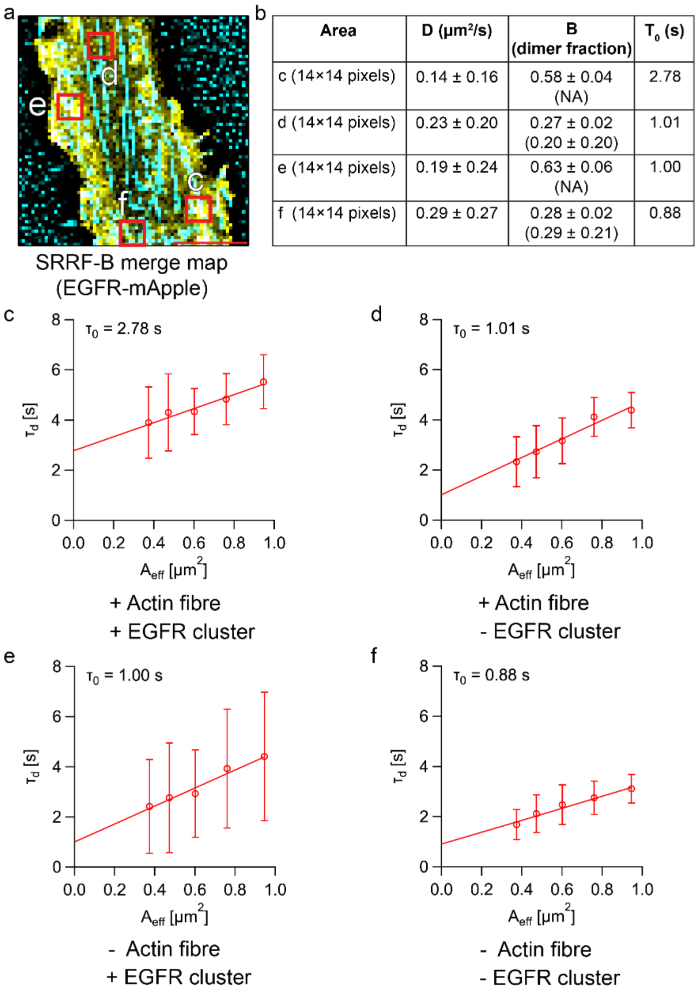


| Area | D (µm²/s) | B (dimer fraction) | $T_0$ (s) |
|---|---|---|---|
| c (14×14 pixels) | 0.14 ± 0.16 | 0.58 ± 0.04 (NA) | 2.78 |
| d (14×14 pixels) | 0.23 ± 0.20 | 0.27 ± 0.02 (0.20 ± 0.20) | 1.01 |
| e (14×14 pixels) | 0.19 ± 0.24 | 0.63 ± 0.06 (NA) | 1.00 |
| f (14×14 pixels) | 0.29 ± 0.27 | 0.28 ± 0.02 (0.29 ± 0.21) | 0.88 |

**Fig. S11: Diffusion law analysis over different areas of a CHO-K1 cell expressing EGFR-mApple cell acquired using a sCMOS: (a)** SRRF-B merge image of a CHO-K1 cell expressing EGFR-mApple. The red boxes (numbered c-f) indicate the different areas tested. **(b)** Table of *D* (mean ± SD), B (mean ± SEM) and intercept values in the chosen areas c-f. The regions with clusters have a higher brightness when compared to regions without clusters. **(c)** Diffusion law in area marked by the red box c. The intercept is 2.78 s. This area (14x14 pixels) was chosen as both an EGFR cluster and an actin fibre were present. **(d)** Diffusion law in area marked by the red box d. The intercept is 1.01 s. This area (14x14 pixels) was chosen as there was an EGFR cluster but no actin fibre present. **(e)** Diffusion law in area marked by the red box e. The intercept is 1.00 s. This area (14x14 pixels) was chosen as there was an actin fibre but no EGFR cluster present. **(f)** Diffusion law in area marked by the red box f. The intercept is 0.88 s. This area (14x14 pixels) was chosen as there was neither an actin fibre nor an EGFR cluster. The scale bar in red measures 5 µm in image (a).

### 4 Appendix

The plasmids used in this study are shown here.

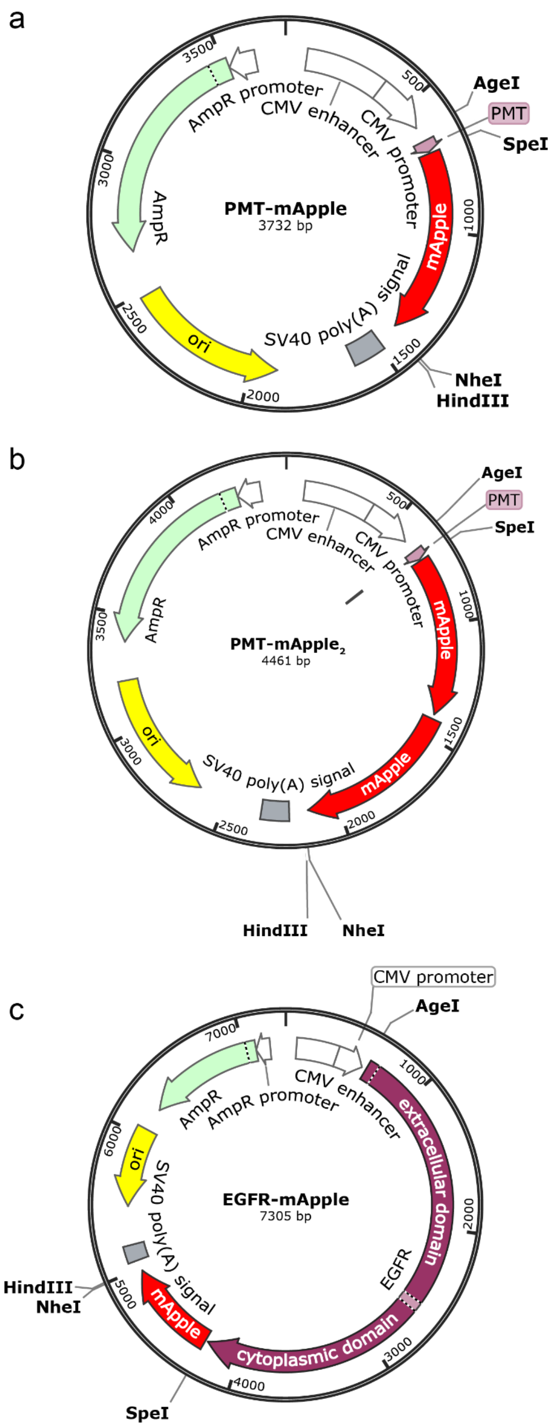


**Fig. S12: Plasmid maps of calibration probes and EGFR-mApple:** Plasmid maps of PMT-mApple, PMT-mApple2, and EGFR-mApple are shown in images **(a)**, **(b),** and **(c)**, respectively. Some of the major sequences are labeled, along with 4 restriction sites – AgeI, SpeI, NheI, and HindIII.
